# Quinoline-based compounds can inhibit diverse enzymes that act on DNA

**DOI:** 10.1101/2024.04.03.587980

**Authors:** Jujun Zhou, Qin Chen, Ren Ren, Jie Yang, Bigang Liu, John R. Horton, Caleb Chang, Chuxuan Li, Leora Maksoud, Yifei Yang, Dante Rotili, Xing Zhang, Robert M. Blumenthal, Taiping Chen, Yang Gao, Sergio Valente, Antonello Mai, Xiaodong Cheng

## Abstract

DNA methylation, as exemplified by cytosine-C5 methylation in mammals and adenine-N6 methylation in bacteria, is a crucial epigenetic mechanism driving numerous vital biological processes. Developing non-nucleoside inhibitors to cause DNA hypomethylation is a high priority, in order to treat a variety of significant medical conditions without the toxicities associated with existing cytidine-based hypomethylating agents. In this study, we have characterized fifteen quinoline-based analogs. Notably, compounds with additions like a methylamine (**9**) or methylpiperazine (**11**) demonstrate similar low micromolar inhibitory potency against both human DNMT1 (which generates C5-methylcytosine) and *Clostridioides difficile* CamA (which generates N6-methyladenine). Structurally, compounds **9** and **11** specifically intercalate into CamA-bound DNA via the minor groove, adjacent to the target adenine, leading to a substantial conformational shift that moves the catalytic domain away from the DNA. This study adds to the limited examples of DNA methyltransferases being inhibited by non-nucleotide compounds through DNA intercalation, following the discovery of dicyanopyridine-based inhibitors for DNMT1. Furthermore, our study shows that some of these quinoline-based analogs inhibit other enzymes that act on DNA, such as polymerases and base excision repair glycosylases. Finally, in cancer cells compound **11** elicits DNA damage response via p53 activation.

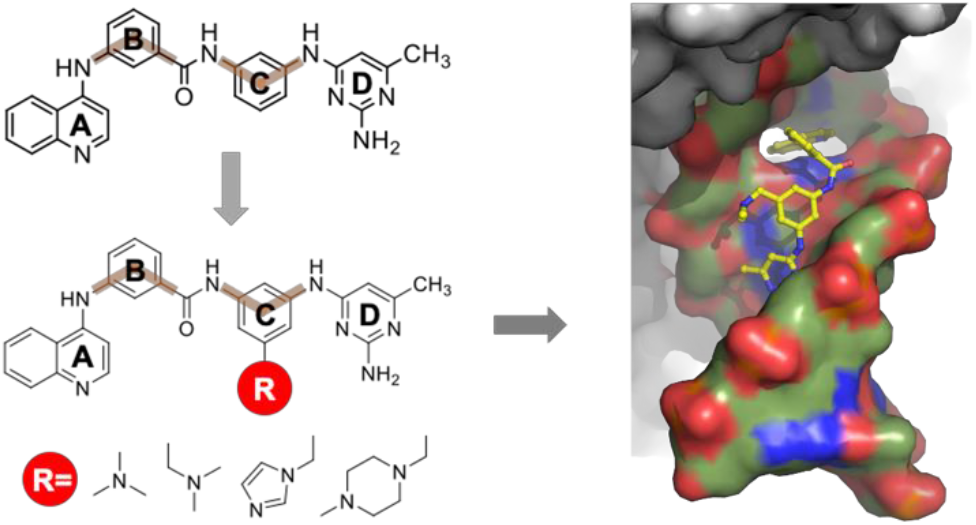

**Highlights:**

- Six of fifteen quinoline-based derivatives demonstrated comparable low micromolar inhibitory effects on human cytosine methyltransferase DNMT1, and the bacterial adenine methyltransferases *Clostridioides difficile* CamA and *Caulobacter crescentus* CcrM.
- Compounds **9** and **11** were found to intercalate into a DNA substrate bound by CamA.
- These quinoline-based derivatives also showed inhibitory activity against various base excision repair DNA glycosylases, and DNA and RNA polymerases.
- Compound **11** provokes DNA damage response via p53 activation in cancer cells.

## Introduction

DNA methylation, mostly in CpG dinucleotides, profoundly impacts chromatin structure and gene expression, affecting inheritance, evolution, and human development and health (Allis et al., 2015; Holliday, 1996). The earliest observations suggesting that epigenetic silencing involved DNA methylation included studies of promoter CpG <u>hyper</u>methylation transcriptionally repressing β-globin genes (van der Ploeg and Flavell, 1980), and tumor-suppressor genes including retinoblastoma (RB) (Greger et al., 1989), von Hippel-Lindau (VHL) (Herman et al., 1994), and p16/CDKN2 (Gonzalez-Zulueta et al., 1995; Herman et al., 1995; Merlo et al., 1995). It has become particularly clear that promoter CpG island hypermethylation of tumor-suppressor genes is a common hallmark of human cancers (Esteller, 2007; Schuebel et al., 2007; Wang et al., 2018). However, DNA <u>hypo</u>methylation is associated with genomic instability, which might be associated with increased tumor heterogeneity (Besselink et al., 2023). Modern whole-genome methylation-based approaches, using cell-free DNA circulating in blood, provide a promising biomarker for early cancer detection (Jamshidi et al., 2022; Luo et al., 2021; Markou et al., 2022).

The two FDA-approved DNA <u>hypo</u>methylating agents, Azacitidine (Vidaza)^1^ and Decitabine (Dacogen)^2^, have been in standard clinical use for some time, particularly in older, medically-challenged patients, for the treatment of certain hematological disorders including myelodysplastic syndromes (Briski et al., 2023; Fenaux et al., 2009; Flotho et al., 2009; Lubbert et al., 2011; Oki et al., 2008; Silverman et al., 2002; Stomper et al., 2019; Yoo and Jones, 2006). However, the dose-limiting toxicity of, and limited patient tolerance for, cytidine analogs, the poor median overall survival for elderly patients with acute myeloid leukemia (<1 year), as well as cytidine analogs’ ineffectiveness in treating solid tumors (Juttermann et al., 1994; Sato et al., 2017; Stomper et al., 2021), have led to a persistent search for non-nucleoside DNMT inhibitors. One such exploration led to the quinoline-based compound SGI-1027 (**1**), which inhibits the activities of CpG-specific methyltransferases (MTases), including mammalian DNMT1, 3A and 3B and bacterial M.SssI (Datta et al., 2009).

Compound **1** contains four ring fragments (A-D in Figure 1A) linked in sequence in a para/para orientation: a 4-aminoquinoline (ring A), a 4-aminobenzoic acid (ring B), a 1,4-phenylenediamine (ring C), and a 2,4-diamino-6-methylpyrimidine (ring D). Rotating each ring fragment’s linkage from the para to the meta or ortho position, or duplicating either the quinoline or pyrimidine moiety, yielded a library of new derivatives (regioisomers and analogs) (Valente et al., 2014). One such analog, MC3343 (**2**), has been further examined *in vitro* and in cancer cells (Cristalli et al., 2022; Manara et al., 2018; Valente et al., 2014; Zwergel et al., 2019). Interestingly, the mechanism of action of compounds **1** and **2** exhibited “a DNA competitive behavior”, “interacted with DNMT only when the DNA duplex was present” or “inhibited DNMTs by interacting with the DNA substrate” (Gros et al., 2015). These biochemical observations led us to re-examine the effects of quinoline-based compounds **1**-**2** and their derivatives (**3**-**17**) on the activities of three human DNMTs yielding 5-methylcytosine (5mC), three bacterial DNA adenine MTases yielding 6-methyladenine (6mA) and a few nucleic acid glycosylases and polymerases.

**Figure 1.**
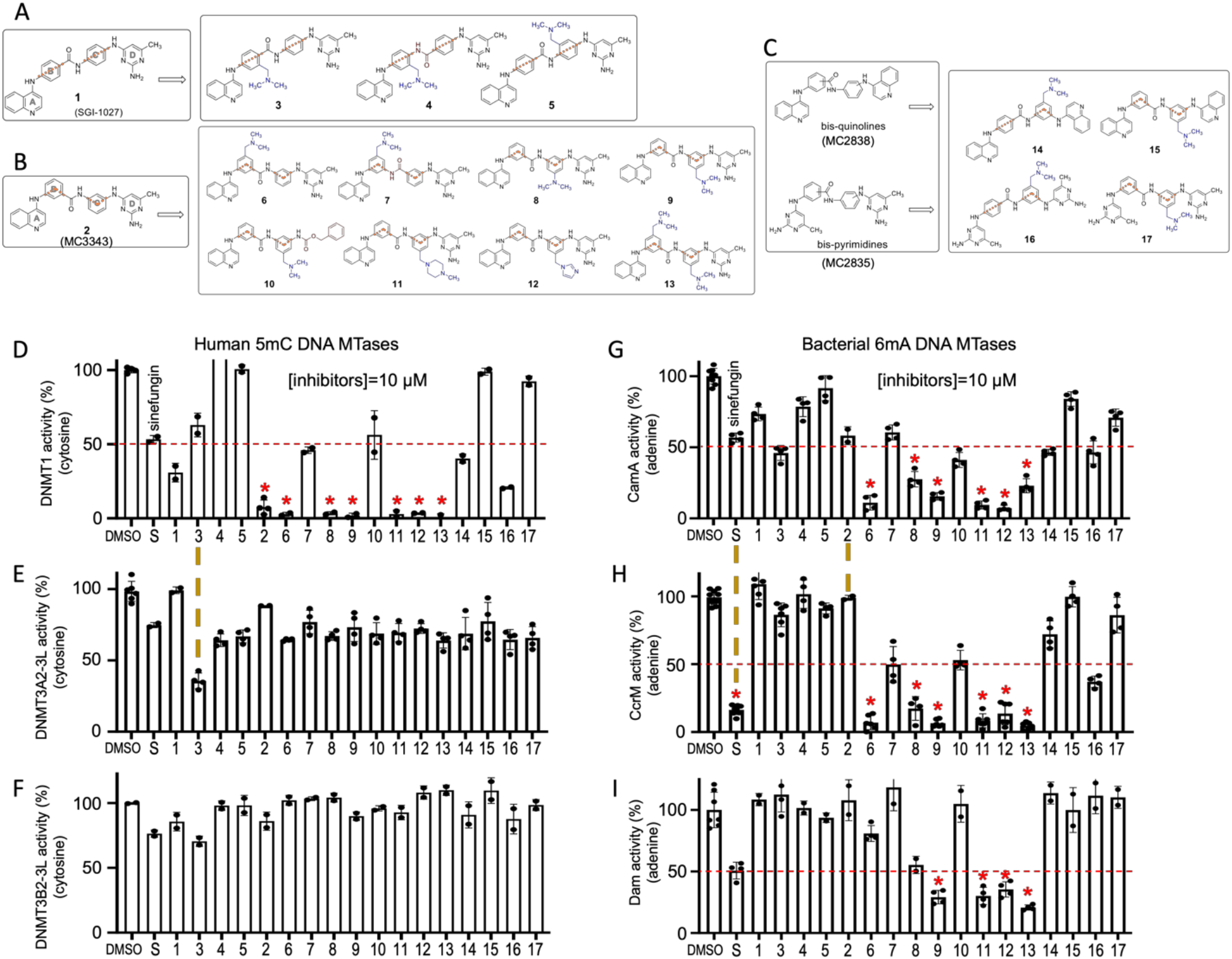
Compounds used in the inhibition study of six MTase activities at a single inhibitor concentration of 10 μM. (**A**) Derivatives (**3-5**) of compound **1** (SGI-1027). (**B**) Derivatives (**6**-**13**) of Compound **2** (MC3343). **(C)** Derivatives (**14**-**17**) of bis-quinolines (MC2838) and bis-pyrimidines (MC2835). The parent compounds MC3343, MC2838 and MC2835 were reported previously as compounds 5, 10 and 11 in (Valente et al., 2014). Link orientations are indicated with orange lines and amino groups are in blue. **(D-F)** Relative Inhibition by compounds against three human 5mC DNA MTases. (G-I) Relative inhibition by compounds against three bacterial 6mA DNA MTases. Compounds indicated by a red asterisk were chosen for further study. DMSO is the control, while S is the known pan-MTase inhibitor sinefungin. The vertical brown dashed lines indicate different inhibition potencies of compound **3** between DNMT1 and DNMT3A/3B, compound **2** between CamA and CcrM/Dam, or sinefungin (S) between CcrM and rest of the five enzymes.

We report here that compounds **9** and **11** exhibit similar low micromolar inhibition potency for both DNMT1 (a 5mC MTase) and CamA (a 6mA MTase). Structurally, compounds **9** and **11** intercalate into CamA-bound DNA, adjacent to the target adenine, via the minor groove. This intercalation leads to a significant conformational shift, moving the catalytic domain away from the DNA. Our findings represent only the second known instance of DNA MTases being inhibited by non-nucleotide compounds through the intercalation of enzyme-bound DNA, following the discovery of dicyanopyridine-based inhibitors against DNMT1 (Horton et al., 2022; Pappalardi et al., 2021).

## Results

### Quinoline-based compounds

In addition to the two parent compounds, **1** and **2**, we prepared fifteen quinoline-based compounds (**3**-**17**), all bearing an amino side chain inserted into one of the phenyl rings of their structures. Compounds **3**-**5** follow the **1** structure (*para*/*para* orientation of the rings B and C) (Figure 1A). Compounds **6**-**13** were built on the structure of **2** (*meta*/*meta* orientation between the rings B and C) (Figure 1B). Compounds **6** and **7** contain a *N,N*-dimethylaminomethyl group attached onto ring B, with either the amide (**6**) or the retroamide (**7**) moiety between the B/C rings. Compounds **8**-**12** have one of the four moieties at ring C: a *N,N*-dimethylamino (**8**), *N,N-*dimethylaminomethyl group (**9**, **10**), (*N*-methylpiperazin-1-yl)methyl (**11**), or a (1*H*-imidazol-1-yl)methyl (**12**) moiety. Differing from **9**, compound **10** replaces the pyrimidine ring (ring D) with a benzyloxycarbonyl moiety. Compound **13** exhibits two *N,N*-dimethylaminomethyl groups inserted respectively at rings B and C. In addition, compounds **14**-**15** and **16**-**17** are derivative of bis-quinolines and bis-pyrimidines respectively (Figure 1C).

### Inhibition of DNA MTases generating 5-methylcytosine or 6-methyladenine

Because the two prototypes **1** and **2** inhibit DNA methylation in a DNA-dependent manner, we tested the new quinoline analogs on three human DNMTs, including the maintenance enzyme DNMT1 acting on a hemi-methylated CpG substrate, and two *de novo* enzymes DNMT3A and DNMT3B acting on unmethylated CpG substrates. Instead of the isolated catalytic MTase domain, we used a construct of DNMT1 expressing residues 351-1,600 (Hashimoto et al., 2012b), which includes the replication foci targeting sequence (RFTS) domain, unmethylated CpG-binding CXXC domain, two copies of the bromo-adjacent homology (BAH) domain and a lysine-glycine (KG)-repeat sequence in addition to the C-terminal catalytic domain (Supplementary Figure S1A). For DNMT3A, we used full-length DNMT3A2, an isoform that lacks the N-terminal region of DNMT3A1 (Chen et al., 2002) (Supplementary Figure S1B), in complex with the effector protein DNMT3L (Jia et al., 2007) (for a review of the different isoforms of human DNMTs, see (Gujar et al., 2019)). Both DNMT3A2 and DNMT3L are normally expressed at high levels in developing germ cells (Sakai et al., 2004). For DNMT3B, we generated an N-terminally truncated DNMT3B2 (Supplementary Figure S1C), equivalent to DNMT3A2, and complexed it with DNMT3L (Zeng et al., 2020). Initially intended as negative (specificity) controls, we also included three bacterial DNA adenine MTases: *Clostridioides difficile* CamA, required for normal sporulation and persistence of infection (Oliveira et al., 2020; Zhou et al., 2021), *Caulobacter crescentus* CcrM, a cell cycle-regulated DNA MTase essential for viability (Horton et al., 2019; Stephens et al., 1996), and *Escherichia coli* DNA adenine MTase (Dam), having roles in coordinating DNA replication initiation, DNA mismatch repair, and the regulation of gene transcription (Horton et al., 2006; Horton et al., 2005; Lobner-Olesen et al., 2005). These various DNA MTases have very little amino acid sequence similarity beyond their catalytic motifs, though their three-dimensional catalytic domain structures are related, and all utilize a base-flipping mechanism to access the target nucleotide (Supplementary Figure S1). Base flipping is often coupled with substantial DNA distortions including bending and unwinding (Cheng and Blumenthal, 2002; Hong and Cheng, 2016; Ren et al., 2022; Roberts and Cheng, 1998).

We conducted an initial inhibition screen, using a compound concentration of 10 μM and employing the MTase-Glo^TM^ biochemical assay (Dong et al., 2020; Hsiao et al., 2016), which detects conversion of the methyl donor *S*-adenosyl-L-methionine (SAM) to *S*-adenosyl-L-homocysteine (SAH). We included sinefungin, a known pan inhibitor of SAM-dependent MTases, as a positive control (Figure 1D-1I). In the case of DNMT1, 10 µM sinefungin and compound **1** demonstrated ∼50% and ∼70% inhibition, respectively (Figure 1D). Of the fifteen new compounds examined, six of them (compounds **6**, **8**, **9**, **11**, **12** and **13**) – all being derivatives of compound **2** – were highly effective on DNMT1 at 10 μM, completely eliminating its activity (Figure 1D). In contrast, the majority of compounds **3**-**17** showed only about 20% inhibition of the DNMT3A2-3L complex, though compound **3** exhibited over 50% inhibition (Figure 1E). Similarly, minimal or no inhibition was observed for DNMT3B2-3L, with the best compound (**3**) demonstrating just ∼20% inhibition (Figure 1F). However, both DNMT3A2-3L and DNMT3B2-3L exhibit low turnover rates under the conditions tested (Supplementary Figure S2A), complicating the detection of changes in enzymatic activity.

Compounds **3**-**17** were tested against three bacterial DNA adenine MTases (CamA, CcrM, and Dam) for their specificity. Unexpectedly, we observed robust inhibition of the activities of the three tested DNA adenine MTases by the same set of six compounds (**6**, **8**, **9**, **11**, **12** and **13**) that had abolished DNMT1 activity (Figure 1G-1I). For comparison, the two parent compounds **1** and **2** exhibited no inhibitory effect at all for CcrM and Dam, and less than 50% inhibition for CamA (Figure 1G). Notably, among these six compounds, compounds **9** and **13** displayed differential activity against CamA, CcrM and Dam, with both compounds showing stronger inhibition of CcrM. Similarly, sinefungin exhibited significant inhibition of CcrM, the strongest potency among the six MTases examined. As sinfungin is a SAM analog, its limited inhibition of CamA might reflect that the enzyme’s unusually weak binding of SAM (Zhou et al., 2021). Preferential inhibition of CcrM by compounds **9** and **13** might reflect the fact that CcrM engages in substantially DNA strand separation, as part of its DNA recognition process, than the other tested MTases (Horton et al., 2019; Reich et al., 2018; Woodcock et al., 2017).

Interestingly, the six most potent compounds (**6**, **8**, **9**, **11**, **12** and **13**) share common chemical features in the backbone, which are reminiscent of the *meta*/*meta* orientation of **2**, maintaining identical relative positions of the ring structures but with modifications in either ring B, ring C, or both (Figure 1B). Changes in the amide fragment from compound **6** to **7** (amide vs inverted amide between B/C rings) or replacement the pyrimidine moiety of ring D from compound **9** to a benzyl carbamate of **10** resulted in loss of inhibition. Conversely, the same modifications on ring B (compound **3**) or ring C (compound **5**) in the backbone of **1** (*para*/*para* orientation) did not yield the same effect. The *bis*-quinoline (**15**) or bis-pyrimidine structure (**17**) resulted in a loss of inhibition ability, highlighting critical importance of the “left” quinoline and the “right” pyrimidine moieties for the derivatives of compound **2**.

### DNMT1 inhibition

We selected the six **2**-mimicries (**6**, **8**, **9**, **11**, **12** and **13**) for a detailed inhibition study, determining their half-maximal inhibitory concentrations (IC_50_) in the presence of 0.1 μM DNMT1, 5.5 μM SAM, and 1 μM DNA substrate (Figure 2A-2F). The six compounds exhibited similar IC_50_ values, ranging from 1.9 to 3.5 μM, with **12** demonstrating 1.8X higher potency than compound **8** (Figure 2G). The six selected quinoline derivatives exhibited undetectable inhibition at low concentrations, transitioning to a steep slope upon reaching the DNA substrate concentration of 1 μM, and plateauing near background levels rapidly thereafter (Figure 2A-2F). This observation suggests that quinoline derivatives operate in a DNA substrate-dependent inhibition mode (Figure 2H-2I).

**Figure 2.**
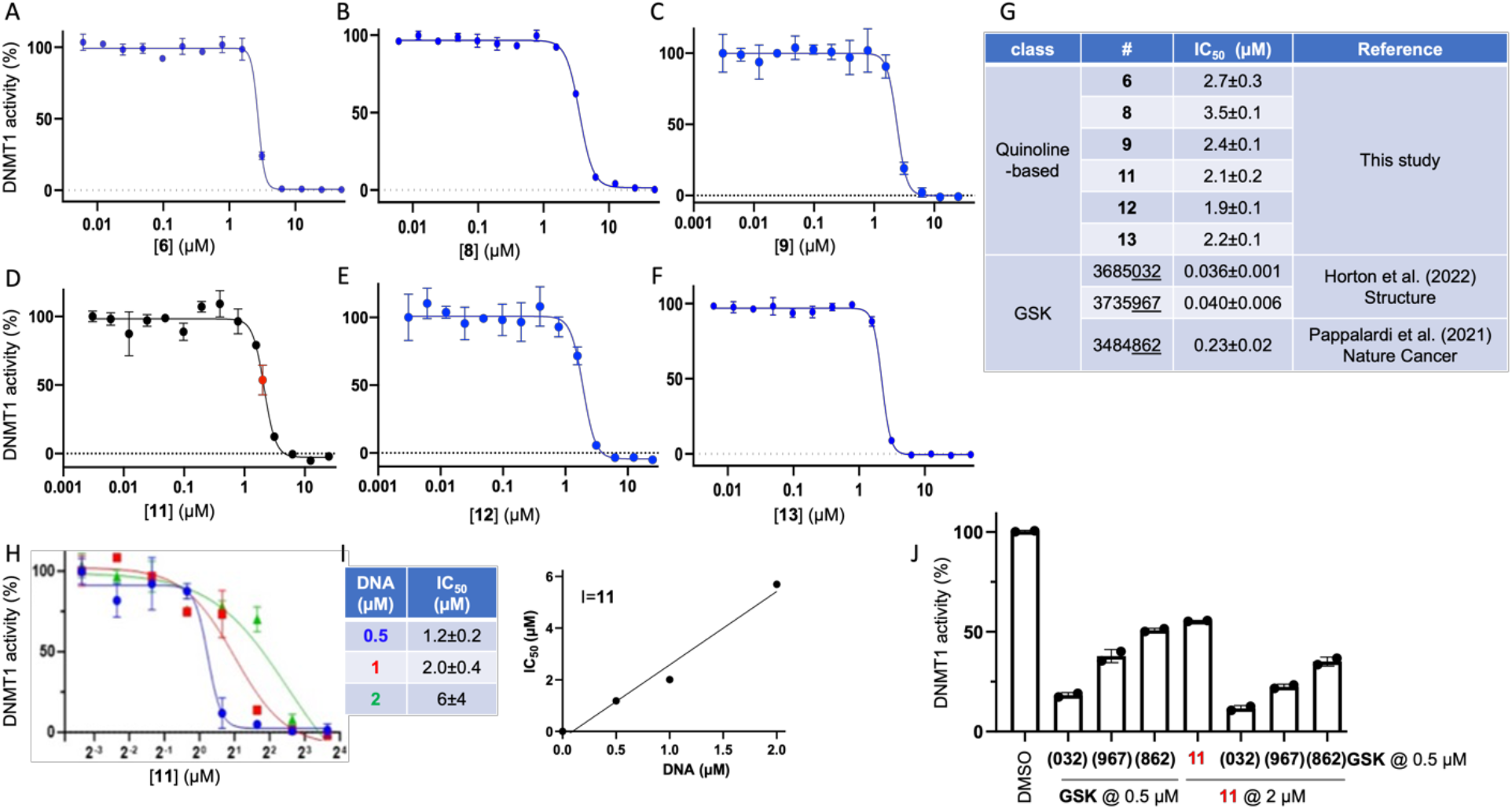
DNMT1 inhibition. (**A-F**) The IC_50_ measurements were made by varying inhibitor concentrations in the presence of 1 μM DNA substrate. (**G**) Summary of the IC_50_ values of inhibition of DNMT1 methylation by quinoline-based compounds and dicyanopyridine-based GSK inhibitors. (**H**) The IC_50_ measurements of compound **11** as a function of substrate DNA concentration. (**I**) Plot of the IC_50_ values of compound **11** verse DNA concentrations. (**J**) Additive effect of inhibition of DNMT1 by GSK inhibitor at 0.5 μM together with compound **11** at 2 μM.

Next, we compared the inhibitory constants with those of three recently developed, dicyanopyridine-containing, non-nucleoside DNMT1-selective GSK inhibitors (Horton et al., 2022; Pappalardi et al., 2021). Under the same reaction conditions, GSK3685032 and GSK3735967 each displayed an IC_50_ value of ∼0.04 μM, while GSK3484862 showed an IC_50_ value of 0.23 µM, making them 10-50 times more potent than the quinoline-based compounds (Figure 2G). At or above their respective IC_50_ values −0.5 μM for GSK inhibitors and 2 μM for compound **11** - the effect of combined treatment is additive, not synergistic (Figure 2J).

We proceeded to examine the impact of the inhibitors in murine embryonic stem cells (mESCs). Unlike other mammalian cells that reply on DNA methylation for viability and proliferation, mESCs can survive without DNA methylation (Chen et al., 2003; Li et al., 1992; Tsumura et al., 2006), making them an ideal cellular system for studying DNA hypomethylation. We used the seven **2**-mimicries (**2**, **6**, **8**, **9**, **11**, **12** and **13**), in conjunction with GSK3685032 and GSK3484862 (Supplementary Figure S3). We assessed global methylation, using methylation of the minor satellite repeats as a proxy, by Southern blot analysis post-digestion with the methylation-sensitive enzyme *Hpa*II (which cuts unmethylated CCGG). After a two-day treatment, only GSK3685032 and GSK3484862 significantly reduced methylation levels, consistent with severe depletion of Dnmt1 (Chen et al., 2023) (Supplementary Figure S3). In contrast, the **2**-related quinoline compounds did not significantly differ from the DMSO control in their effect on methylation, despite their strong *in vitro* inhibition of DNMT1 (Figure 2). This discrepancy may be partly due to their pharmacokinetic properties, including their capacities to enter the cells and the nuclei, where DNA methylation occurs, their stability in cells, or their interactions with various other enzymes (see below).

### CamA inhibition

We determined the IC_50_ values, which ranged from 2-4 μM, for the four quinoline compounds (**6**, **9**, **11** and **12**) (Figure 3A) that had exhibited the most potent inhibition against CamA in a single-dose experiment (Figure 1G). These values are in a similar range as we observed for DNMT1 inhibition. Compound **12** inhibited both DNMT1 and CamA with comparable potency (IC_50_ ∼ 2 μM). Subsequently we compared these results to MC4741, a recently-developed selective CamA inhibitor that features an adenosine analog carrying a 3-phenylpropyl moiety attached to the adenosyl N6 atom (Zhou et al., 2023). MC4741 exhibited stronger inhibition, with an IC_50_ value of 0.7 μM (Figure 3B), 3X lower than that of compound **12** and 5-6X lower than that of the other three quinoline compounds (Figure 3C). When used in combination, at their respective IC_50_ values, compounds **9** (or **11**) together with MC4741 exhibited additive effect (Figure 3D).

**Figure 3.**
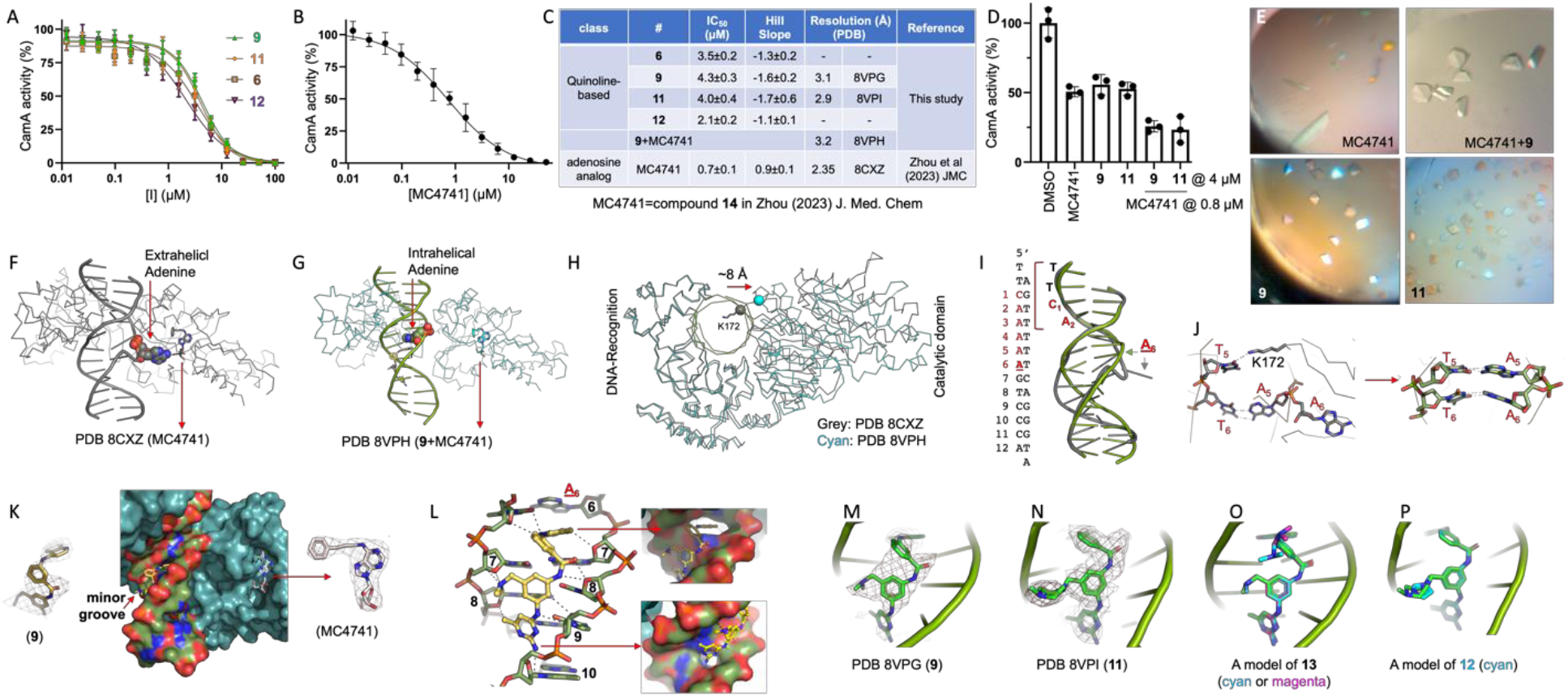
CamA inhibition. (**A**) The IC_50_ measurements were made by varying inhibitor concentrations of four quinoline-based derivatives. (**B**) The adenosyl analog MC4741 inhibits CamA activity. (**C**) Summary of the IC_50_ values of inhibition of CamA methylation by various inhibitors, and X-ray information (PDB accession numbers and corresponding resolutions). (**D**) Additive effect of inhibition of CamA by two quinoline-based compounds (**9** or **11**) at 4 μM together with compound MC4741 at 0.8 μM. (**E**) Examples of CamA-DNA crystal morphology. (**F**) The target adenine (space-filled) projects out of the DNA double helix even when the SAM binding site is occupied by an analog like MC4741 (PDB 8CXZ). (**G**) The target (methylatable) adenine remains intrahelical with the introduction of compound **9**. (**H**) Upon binding of compound **9**, the catalytic domain of CamA pulls back from the DNA, but the DNA-recognition domain of CamA remains engaged with the DNA. The small spheres indicate the Cα positions of Lys172. (**I**) The interactions remain the same with the sequence 5’ to the target adenine. (**J**) In the flipped-out adenine structure (PDB 8CXZ), DNA has substantial distortions including a base pair rearrangement (A5:T6) and a gained protein side chain interaction (Lys172) with the orphaned base T5 (*left* panel). In the structure where the target adenine remains intrahelical (PDB 8VPH), the base pairing has been restored for two adjacent AT base pairs at positions 5 and 6 (*right* panel). (**K**) Surface representation of ternary complex of CamA-DNA-**9**-MC4741. The omit electron densities are shown, contoured at 4α above the mean, for MC4741 and compound **9**, respectively. (**L**) Compound **9** engages in stacking and van der Waals interactions with CamA-bound DNA in the minor groove. Two widened helical rises along the DNA axis correspond to the two ends of compound **9**. (**M**) Compound **9** bound with CamA-DNA (PDB 8VPG). (**N**) Compound **11** bound with CamA-DNA (PDB 8VPI). (**O**) A model of compound **13**. (**P**) A model of compound **12**.

MC4741 inhibits CamA activity by occupying the binding pocket of methyl donor SAM (Zhou et al., 2023). To clarify the inhibition mode of the new quinoline derivatives, we meticulously explored co-crystallization conditions for CamA-DNA complexes in the presence of compound. We determined three structures of CamA-DNA complexes with compounds **9** or **11** individually, or compound **9** together with MC4741, in a resolution range of 2.9-3.2 Å (Figure 3C). The introduction of the quinoline derivatives into the crystallization mix altered the crystal shapes from elongated rods to diamonds (Figure 3E), though the crystals were formed in the same crystallographic space group and with similar unit cell dimensions (Table S1). Within the crystallographic asymmetric unit, there are three CamA-DNA complexes, and only one of the three complexes showed a bound inhibitor (**9** or **11**). We made the following observations from the quinoline-bound complexes.

CamA shares a common feature with other structurally-characterized DNA MTases acting on cytosine and adenine (Roberts and Cheng, 1998; Woodcock et al., 2020). Specifically, like all other known DNA MTases, CamA flips its target adenine out of the DNA double helix (Zhou et al., 2021). This action occurs even when the SAM binding site is occupied by an analog like MC4741 (Figure 3F). However, with the introduction of compound **9** or **11**, the target adenine either doesn’t flip or it returns to its normal position within the double helix (Figure 3G). Superimposition of the complexes reveals that, upon binding of compound **9**, the DNA-recognition domain of CamA remains engaged with DNA, but the catalytic domain retracts by approximately 8 Å (Figure 3H). More specifically, the DNA-recognition domain stays in close contact with the five base-pairs of the recognition sequence 5’ to the target adenine in the major-groove, such that this portion of the DNA is undisturbed (Figure 3I). However, the catalytic domain retraction disrupts the interaction between the enzyme’s catalytic residues and the DNA, specially the Lys172-base T5 interaction, and restores the base pairing of two adjacent AT base pairs at positions 5 and 6 (Figure 3J). The Lys172-containing active-site loop, which normally penetrates the DNA minor groove, undergoes large movement and becomes less ordered when the enzyme transitions in the presence of compound **9**.

Compound **9** engages with DNA in the minor groove (Figure 3K). The aminoquinoline moiety, intercalates itself into the DNA right after the base pair that includes the target adenine. This insertion occurs chiefly through stacking interactions with the base pairs at positions 6 and 7 (Figure 3L). The subsequent molecular segments, 3-aminobenzoic acid and 1,3-phenylenediamine, extend across the subsequent three base pairs (7 to 9). The terminal component, a diamino-6-methylpyrimidine moiety, wedges in between the base pairs at positions 9 and 10 (Figure 3L). Beyond the stacking provided by the aminoquinoline, compound **9**’s primary interactions with the DNA are van der Waals forces with six deoxyribose rings, three on each strand. *N,N*-dimethylaminomethyl addition to the 1,3-phenylenediamine moiety allows two van der Waals contacts with the deoxyribose of cytosine at position 7. Additionally, the amino and methyl groups on the diamino-6-methylpyrimidine moiety form a hydrogen bond with the cytosine O2 atom at position 10, and a van der Waals interaction with the deoxyribose O4 atom of adenine at position 8, respectively (Figure 3L). The helical rise along the DNA axis is extended at two specific points, corresponding to the two ends of compound **9**: between base pairs 6 and 7 due to the intercalation of the aminoquinoline moiety, and between base pairs 9 and 10 where the diamino-6-methylpyrimidine is positioned (Figure 3L).

In addition, compounds **9** and **11** exhibited similar binding modes when tested independently, without the presence of MC4741 (Figure 4M and 4N). Building upon the structure of compound **9**, we introduced an additional *N,N*-dimethylaminomethyl group onto ring B, creating compound **13** (Figure 3O). In this modeled structure, the second *N,N*-dimethylaminomethyl group of compound **13** is oriented away from the DNA, facing towards the solvent. For compound **11**, we replaced its *N*-methylpiperazin-1-yl moiety with an imidazole ring to form compound **12** (Figure 3P). This substitution of a positively charged imidazole ring allows potential interaction with the nearest DNA backbone phosphate group, potentially contributing to compound **12**’s twofold increased potency, as reflected in its IC_50_ value.

**Figure 4.**
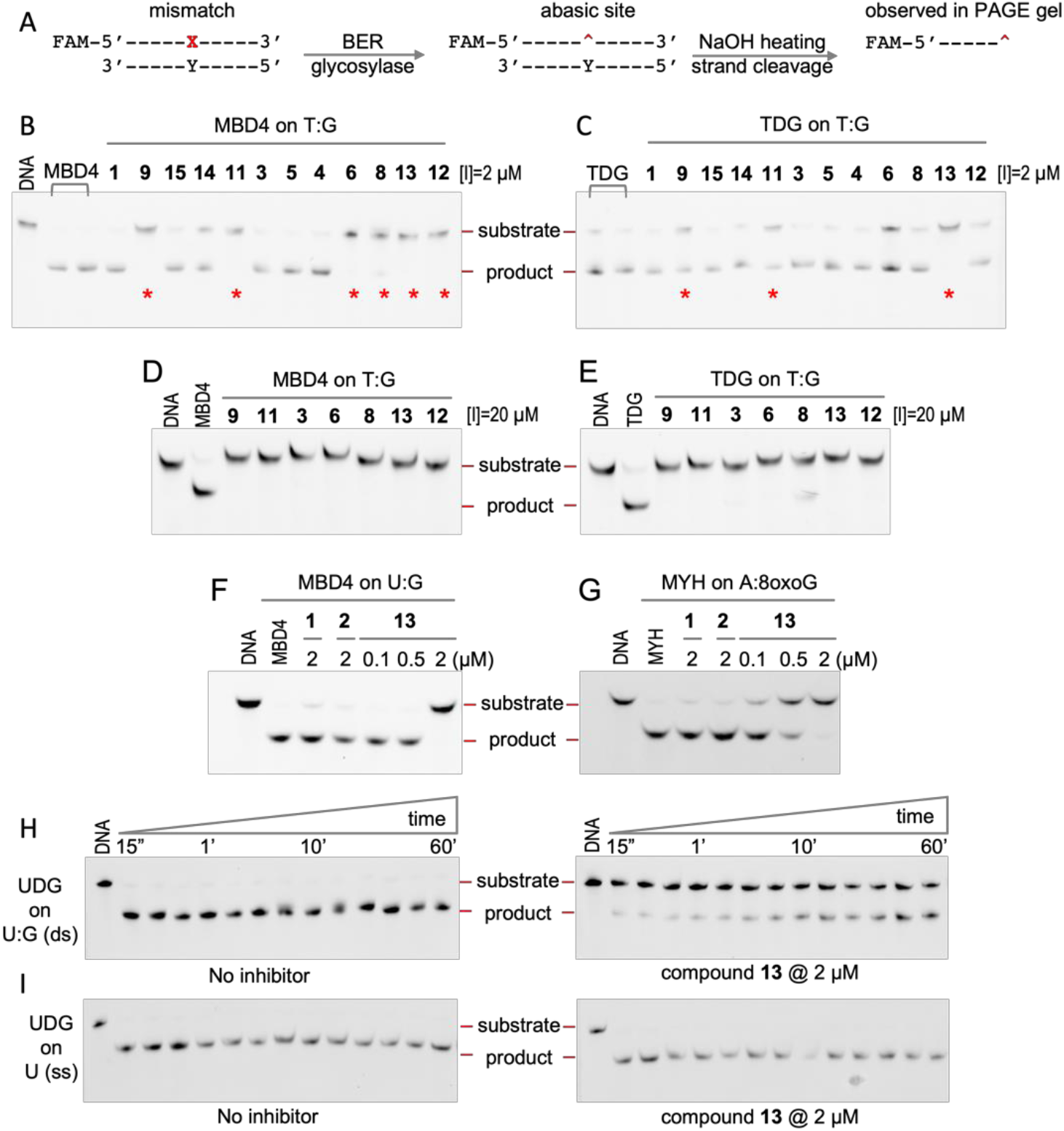
Inhibition of base excision repair (BER) glycosylases on dsDNA containing mismatches. (**A**) Schematic of strand cleavage at an abasic site in dsDNA arising from glycosylase-induced and NaOH-treated. (**A-E**) Inhibition of MBD4 and TDG activities on T:G mismatches at inhibitor concentrations of 2 μM (**B-C**) or 20 μM (**D-E**). (**F**) Inhibition of MBD4 activity on U:G mismatch. (**G**) Inhibition of MYH activity on adenine opposite to 8-oxoguanine. (**H**) Activity of UDG on uracil of dsDNA without (left panel) and with compound **13** at 2 μM (right panel). (**I**) Activities of UDG on uracil of ssDNA without (left panel) and with compound **13** at 2 μM (right panel).

The insertion of planar moieties between adjacent bases is a common mechanism for drug intercalation into DNA or RNA (Satange et al., 2019; Satpathi et al., 2021; Soni et al., 2017). However, the unique aspect here is the specific insertion of aminoquinoline-based derivatives after the target adenine, within the sequence bound by CamA. While there is no observed direct interaction between the compound and CamA in the current structure, it seems likely that the inhibitor competes with or displaces the enzyme’s active-site loop from the DNA’s minor groove. This action results in the reestablishment of Watson-Crick base pairing while the DNA remains bound by the enzyme. In addition, the quinoline-based compounds (**9** or **11**) do not interact with DNA or CamA independently, but reduce CamA-DNA interaction (Supplementary Figure S4). This observation suggests that the quinoline-based compounds specifically alter the enzyme’s interaction with DNA, providing a structural basis for its inhibitory effect.

### Inhibition of base excision repair DNA glycosylases

The finding that compounds **9** and **11** intercalate into CamA-bound DNA, reversing base flipping, prompted us to explore whether our quinoline derivatives can inhibit other DNA base-flipping enzymes. Specifically, DNA <u>b</u>ase <u>e</u>xcision repair (BER) enzymes such as uracil-DNA glycosylase (UDG), thymine-DNA glycosylase (TDG), methyl-CpG-binding domain protein 4 (MBD4) and MutY homolog (MYH) utilize a base-flipping mechanism for excising bases (Hashimoto et al., 2012a; Hashimoto et al., 2012c; Slupphaug et al., 1996). Initially, we assayed MBD4 and TDG base excision activities on a T:G mismatch substrate (with the mispaired T-strand being FAM-labeled in Figure 4A), using an inhibitor concentration of 2 μM. Similar to their effect on DNMT1 and CamA, the same six compounds (**6**, **8**, **9**, **11, 12** and **13**) inhibited MBD4 activity (Figure 4B). However, only compound **13** almost completely inhibited TDG activity, while the other compounds exhibited lesser inhibition (Figure 4C). The variance in inhibition levels between MBD4 and TDG was not observed when the inhibitor dose was increased tenfold (Figure 4D, E). In addition, using compound **13** as an example, we noted that this compound does not independently bind with mismatched DNA (Supplementary Figure S5). The inhibition effect on MBD4 is not substrate specific, as compound **13** inhibits MBD4 activity on a U:G mismatch (Figure 4F). Moreover, compound **13** also effectively inhibits the excision activity MYH in the removal of adenine opposite 8-oxo-2’-deoxyguanosine (8-oxoG) (Figure 4G). Notably, the two progenitor compounds, **1** and **2**, do not exhibit effective inhibition at a concentration of 2 μM (Figure 4B, C, F and G). The lack of effective inhibition by the original compounds at this concentration underscores the importance of molecular modifications in enhancing or altering the inhibitory capabilities of these compounds against base-flipping enzymes.

Continuing our investigation, we explored the excision of uracil by UDG, an enzyme notable for its activity on both double-strand (ds) and single-strand (ss) DNA. We found that compound **13** exhibited less potent inhibition of UDG activity on dsDNA substrates (Figure 4H) compared to the other glycosylases examined in this study (MBD4, TDG, and MYH). However, a critical observation is that compound **13** does not inhibit UDG activity on ssDNA (Figure 4I). This differential effect highlights the substrate specificity of these quinoline-based derivatives using a mechanism of intercalation with dsDNA, as demonstrated by CamA (Figure 3).

### Inhibition of polymerases

Next, we considered DNA polymerases, such as Y-family translesion DNA polymerase η (Pol η) and A-family repair DNA polymerase θ (Pol θ), which are considered potential targets for cancer therapy (Lange et al., 2011; Tsegay et al., 2023). Using sinefungin, a known pan MTase inhibitor, as a control, which as expected showed no impact on polymerase activity (Figure 5A and Supplementary Figure S6), we found that compounds **9** and **13** inhibited >50% of Pol η’s activity at a compound concentration of 10 µM (Figure 5A). Further, in the presence of 50 µM deoxynucleoside triphosphate substrates (dNTP), compounds **9** and **13** exhibited IC_50_ values of 7.5 µM and 3.9 µM, respectively, in inhibiting Pol η activity (Figure 5B, C).

**Figure 5.**
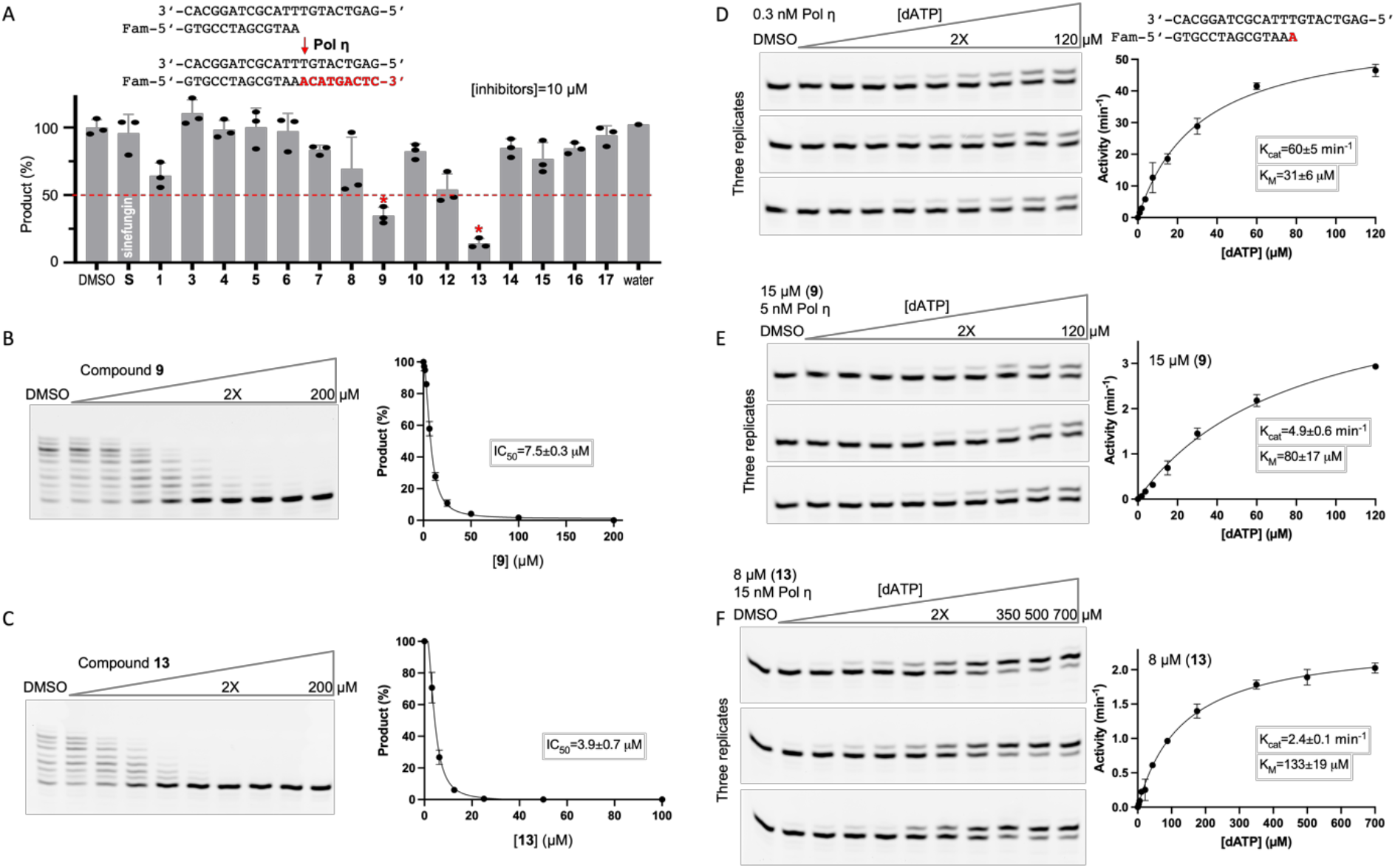
Inhibition of DNA Pol η. (**A**) The DNA synthesis reaction uses DNA template (top strand) and 5’-fluorescein-labeled primer (bottom strand). Relative inhibition of pol η at a single inhibitor concentration of 10 μM. [S, sinefungin; compounds **2** and **11** were unavailable at the time of these assays.] (**B-C**) Inhibition of Pol η activity with increased concentrations of compounds **9** (panel B) or **13** (panel C). (**D-F**) Kinetics of Pol η on the first adenine incorporation in the absence of inhibitor (panel D), presence of compound **9** (panel E), or of compound **13** (panel F).

We then conducted steady-state kinetic assays focusing on the first incoming nucleotide incorporation (dATP) (Figure 5D). Under the laboratory conditions (Methods), Pol η exhibited a catalytic rate constant (*k*_cat_) of 60 min^−1^ and a Michaelis constant (K_M_ of dATP) of 31 µM (Figure 5D). When exposed to a concentration of compound **9** or **13** at twice their respective IC_50_ values, we observed significant changes in the enzyme kinetics: a 12-fold or 25-fold reduced *k*cat values for compounds **9** (5.0 min^−1^) and **13** (2.4 min^−1^) respectively, and a 2-4 fold increase in K_M_^dATP^ values (Figure 5E, F). This translates to a 30-fold or 95-fold reduction in catalytic efficiency for compounds **9** and **13**, respectively, as evidenced by comparing the *k*_cat_/K_M_^dATP^ value of 1.9 min^−1^µM^−1^ (with no inhibition) to 0.06 min^−1^µM^−1^ (compound **9**) or 0.02 min^−1^µM^−1^ (compound **13**).

Using the same conditions, we observed that at a concentration of 10 µM, none of the compounds inhibited Pol θ activity by more than 20% (Supplementary Figure S6A). However, at a fivefold higher concentration (50 µM), compounds **1** and **9** achieved ∼50% inhibition, while compound **13** reached approximately 75% inhibition. Notably, compounds **9** and **13** demonstrated higher IC_50_ values of 68 µM and 14 µM, respectively (Supplementary Figure S6B-C), making them 9 times and 3.6 times less effective as inhibitors of Pol θ compared to Pol η. The basis for differential inhibition of these two repair polymerases, belonging to different families with distinct difference sequences and structures, remains elusive, but the important point here is that these DNA-intercalating inhibitors of DNA MTases also inhibit some DNA polymerases.

We further evaluated the quinoline derivatives against HIV reverse transcriptase (RT), an enzyme that synthesizes DNA on primed RNA templates. Remarkably, nearly all of the compounds showed approximately 50% inhibition at a concentration of 10 µM (Supplementary Figure S6D). When tested at a higher concentration of 50 µM, four of these compounds (**3**, **7**, **9**, **12**) abolished the activity of HIV RT to undetectable level (Supplementary Figure S6D).

While our focus here is on enzymes that act on DNA, in our final set of experiments, we tested the quinoline derivatives against a poliovirus RNA-dependent RNA polymerase (RdRp), which synthesizes RNA on primed RNA templates. Intriguingly, the parent compound **1** exhibited the most significant inhibition, with around 75% effectiveness at a concentration of 10 µM (Supplementary Figure S6E). Notably, the RdRp enzyme demonstrated a unique response compared to the other tested polymerases and MTases, particularly in its sensitivity to compound **1**. This finding suggests that quinoline derivatives might differentiate between RNA and DNA duplexes in their mechanism of action. However, again, the key point for the purposes of this study is that these target-specific intercalating drugs inhibit a wide range of enzymes that act on nucleic acids.

### Compound 11 elicits DNA damage response via p53 activation

Given the in vitro inhibitory activity of quinoline-based derivatives on various enzymes acting on DNA substrates, we investigated the effects of compound **11** on the viability of A549 human non-small cell lung carcinoma (NSCLC) cells. Through luminescence-based cell viability assays, we observed that treatment with **11** at a concentration of 4 µM for three days resulted in a reduction of A549 cell growth to ∼50% compared to DMSO-treated control cells (Figure 6A). In contrast, cells exposed to GSK3484862 exhibited growth rates similar to control cells, as we had observed previously (Chen et al., 2023), whereas GSK3685032 displayed higher toxicity than GSK3484862 but was less effective in reducing cell viability than compound **11** (Figure 6A). To minimize the impact of cell death on subsequent analyses, A549 cells were treated for one day in the following experiments.

**Figure 6.**
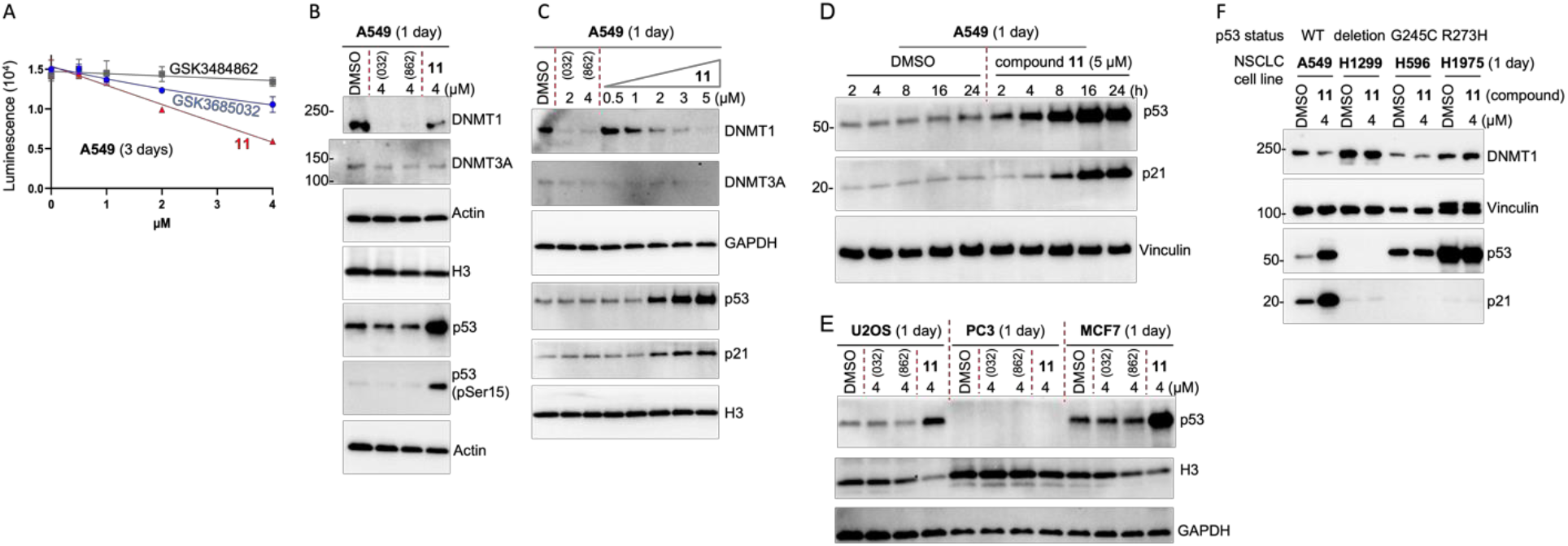
The effect of compound 11 in A549 cells. **(A**) Cell viability (relative to DMSO treatment). N = 3 independent experiments with triplicates (mean ± SD). (**B**) Western blots showing endogenous levels of DNMT1, DNMT3A, p53 and pSer15 of p53 following treatment with 4 µM of GSK3484<u>862</u>, GSK3685<u>032</u> and compound **11** for 24 h. (**C**) Dose-dependent decrease of DNMT1 and increase of p53 and p21 by compound **11** treatment in A549 cells. **(D)** Time-dependent increase of p53 and p21 by compound **11** treatment in A549 cells. (**E**) Western blot confirms increased p53 levels in osteosarcoma U2OS (left) and breast-cancer derived MCF7 cells (right), but not in prostate cancer-derived PC3 (middle). (**F**) Treatment of compound **11** in a set of non-small cell lung cancer (NSCLC) cell lines, encompassing a variety of p53 genetic backgrounds.

Recent studies have highlighted the efficacy of dicyanopyridine-based GSK inhibitors in degrading DNMT1 (Chen et al., 2023). We reproduced this observation, demonstrating a complete depletion of DNMT1 in A549 cells treated with GSK inhibitors (Figure 6B). Interestingly, similar treatment conditions with compound **11**—a 4 µM concentration for one-day—resulted in a decreased DNMT1 protein level, albeit to a lesser extent (Figure 6B). We observed no significant changes in DNMT3A protein levels. The most notable distinction between these two classes of inhibitors lies in their impact on p53—a protein crucial for the cellular DNA damage response (Abuetabh et al., 2022; Steffens Reinhardt et al., 2023)—and the phosphorylation of p53 at serine 15 (Ser15). This post-translational modification is key for the activation and stabilization of p53 following DNA damage (Siliciano et al., 1997), highlighting a potential differential mechanism of action between the two chemotypes. Moreover, treatment of A549 cells with compound **11** led to dose- and time-dependent increases in p53 and its downstream regulator p21 (Figure 6C, D). This observation supports compound **11** having stronger cellular toxicity (via a DNA-damage effect) than the GSK compounds (Figure 6A).

Additionally, we confirmed that compound **11** treatment elevated p53 protein levels in osteosarcoma-derived U2OS cells and in estrogen receptor-positive, breast cancer-derived MCF7 cells (Figure 6E). As a control, p53 protein was undetectable in prostate cancer-derived, androgen-insensitive PC3 cells, due to deletion of the *TP53* gene (Carroll et al., 1993; Isaacs et al., 1991; Rubin et al., 1991).

Next, we extended our investigation of compound **11**’s effects to a broader array of NSCLC cell lines, encompassing a variety of p53 genetic backgrounds: wild-type p53 (A549), p53 deletion (NCI-H1299), and p53 hotspot mutations Gly245-to-Cys (NCI-H596) and Arg273-to-His (NCI-H1975) (Figure 6F). Treatment of the p53 wild-type (A549) cells led to a partial reduction in DNMT1 levels and the activation of p53 and p21, an indication of DNA damage/cellular stress caused by the compound treatment. However, no significant impact on p53/p21 level was observed in cell lines with p53 deletion or those harboring p53 mutations.

Unlike the cancer cell lines studied here, mouse embryonic stem cells (mESCs) exhibited no alteration in p53 levels following exposure to compound **11** and related quinoline derivatives (Supplementary Figure S3). mESCs are characterized by rapid cell cycling and a notably brief G1 phase. The role of p53 in mESCs, particularly its involvement in the DNA damage response, is multifaceted and influenced by various factors, including culture conditions and cell density (Ayaz et al., 2022). The question of whether these quinoline compounds are less cytotoxic or if the cytotoxicity they induce elicits a minimal p53 response in mESCs has yet to be resolved.

## Discussion

### DNA intercalating agents as DNMT inhibitors

Recently, we investigated the structural and biochemical interactions of human DNMT1 with dicyanopyridine-based inhibitors (Horton et al., 2022; Pappalardi et al., 2021). These inhibitors are unique in their chemotype, featuring a planar dicyanopyridine core that specifically intercalates into DNMT1-bound DNA substrate (Figure 7A). Our structural insights into these dicyanopyridine-based compounds, combined with biochemical data showing DNA-dependent inhibition by quinoline derivatives (Gros et al., 2015), prompted us to re-examine the mechanism of the quinoline derivatives in inhibiting two distinct classes of DNA MTases – those generating 5mC and those generating 6mA.

**Figure 7.**
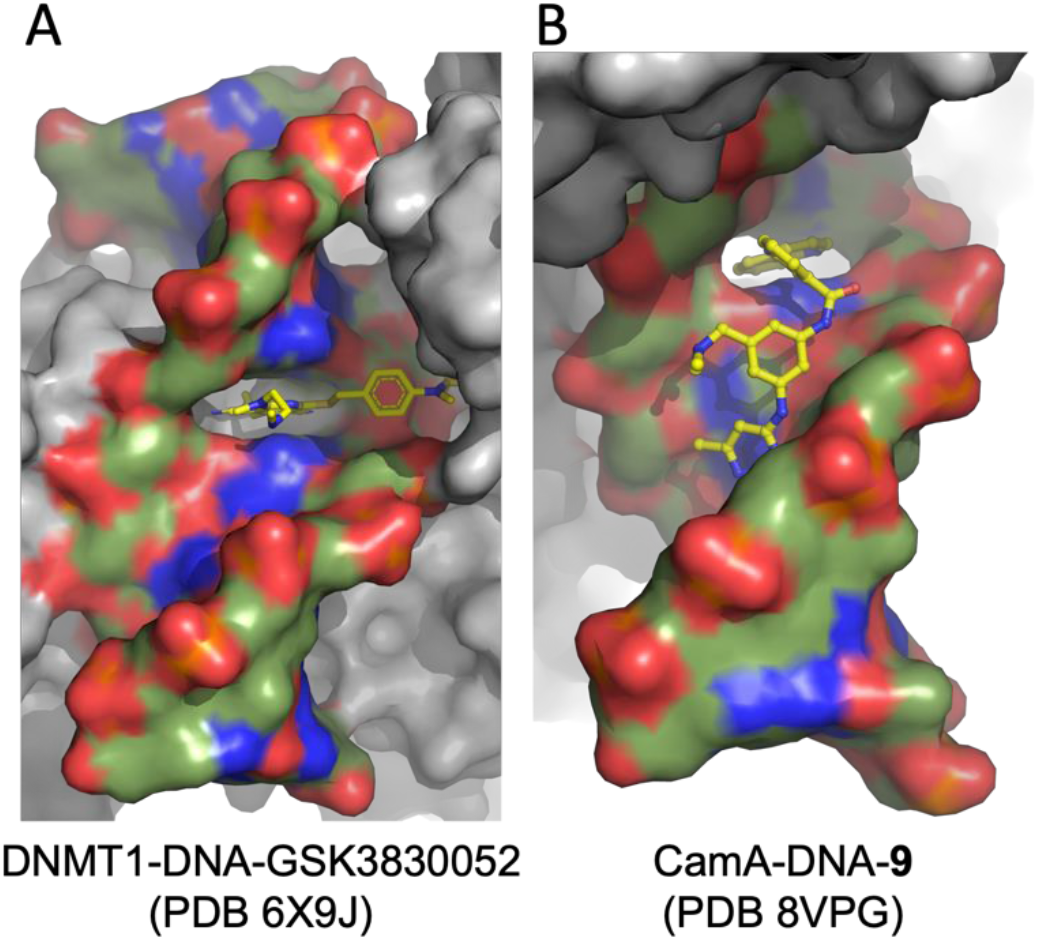
Two examples of DNA intercalating agents as DNA MTase inhibitors. Surface representation of protein (gray) – DNA (blue for nitrogen atoms, red for oxygen atoms, and green for carbon atoms) – compound (yellow sticks) ternary complexes. (**A**) Dicyanopyridine-based GSK3830052 intercalates into DNMT1-bound DNA (PDB 6X9J). (**B**) Quinoline-based compound **9** intercalates into CamA-bound DNA (PDB 8VPG).

We identified three key similarities between dicyanopyridine-based and quinoline-based inhibitors, applicable to both compound types. (1) Intercalation into DNA – both dicyanopyridine and aminoquinoline moieties intercalate into the DNA at sites where it is bound by MTases (Figure 7), disrupting the normal DNA helical rise. (2) Conformational shift in MTase catalytic domain – the intercalation induces a significant conformational shift of the MTase catalytic domain, moving away from the specific DNA sequence, but does not depend on catalytic competence at least in the case of DNMT1: an active-site cysteine-to-serine substitution does not affect the dicyanopyridine compound interaction with DNA in the presence of the mutant DNMT1 (Chen et al., 2023). (3) Restoration of base pairing – the interaction between the enzyme’s active-site residues and the extrahelical target nucleotide (cytosine or adenine) is disrupted, allowing these flipped-out bases to return to their normal intrahelical positions. While the initial mechanism directing these inhibitors to their specific binding sites remains unclear, given that they bind to neither the DNA alone nor the enzyme alone, we hypothesize that the DNA deformation caused by the base-flipping action of MTases might create an entry point for these intercalating agents. Supporting this hypothesis, we also observed that these compounds inhibit DNA base excision repair glycosylases such as MBD4, TDG, MYH and UDG (Figure 4), which employ a base-flipping mechanism for excising bases (Hashimoto et al., 2012a; Hashimoto et al., 2012c; Slupphaug et al., 1996). This led us to speculate that quinoline-based analogs may have a broad spectrum of activity against various DNA-acting enzymes, such as polymerases, that contain a junction of dsDNA and ssDNA during catalysis.

### Compound selectivity vs. potency

A significant distinction between dicyanopyridine-based and the quinoline-based inhibitors studied here is their selectivity. Dicyanopyridine-based inhibitors tested to date exhibit exclusive specificity towards DNMT1, whereas quinoline-based inhibitors can inhibit both DNMT1 as well as the three adenine DNA MTases, and four BER glycosylases in our tests. This difference in selectivity is thought to arise from the interaction of dicyanopyridine-based inhibitors with the active-site loop of DNMT1 (Figure 7A) (Horton et al., 2022; Pappalardi et al., 2021), which is likely crucial for their DNMT1-specific action. In contrast, our structural analyses, representing only a snapshot along the inhibition pathway, did not reveal a direct interaction between the quinoline-based compounds **9** and **11** and CamA (Figure 7B). This absence of direct interaction in the observed CamA structures suggests that quinoline-based compounds might have a wider range of targets, potentially inhibiting various other MTases or enzymes that interact with nucleic acids. Aligned with this concept, the quinoline-based compounds demonstrate more pronounced cytotoxic effects than dicyanopyridine-based DNMT1 inhibitors.

However, not all quinoline-based inhibitors exhibited uniform behavior. For instance, the parent compound **2** displays a distinct selectivity pattern, with 10 µM eliminating nearly all DNMT1 activity, inhibiting CamA by about 50%, and showing no inhibitory effect on CcrM or Dam (Figure 1). However, derivatives of **2**, such as compounds **9** and **11**, have higher inhibitory potency across all four enzymes and lack obvious selectivity among the four.

Regarding polymerases, compounds **1** and **13** demonstrated contrasting inhibitory effects on DNA repair polymerase Pol η and the RNA-dependent RNA polymerase RdRp. Compound **13** is most effective against Pol η, while compound **1** shows more than 50% inhibition at 10 µM. Conversely, compound **1** is the most potent against RdRp, whereas compound **13** does not detectably inhibit it, even at a concentration of 50 µM (Supplementary Figure S6E). These observations highlight the potential for fine-tuning the potency and selectivity of quinoline-based inhibitors to target specific enzymes more effectively.

### Limitations of the study

This study acknowledges several limitations, particularly in the exploration of DNA and RNA interactions with proteins and enzymes acting on double-stranded substrates. Such interactions often result in significant structural changes to the nucleic acids, including kinking, bending, unwinding, base flipping, and separation of strands, which may facilitate the binding of small molecule intercalators. Our investigation was limited to a select few of these enzymes, leaving a vast array of potential targets yet to be explored. Moreover, the demonstrated specificity of compound **13** in inhibiting UDG activity on dsDNA over ssDNA highlights the possibility of developing inhibitors that are substrate selective for enzymes acting on both forms of substrates, though it also underscores the need for extensive research to fully understand and exploit these mechanisms.

### Significance

Our research revealed that treating various cancer cells with compound **11**, particularly those with wild-type p53, significantly elevates the protein levels of the tumor suppressor p53 and its downstream effector p21. Once activated, p53 can trigger cell cycle arrest, apoptosis (programmed cell death), and senescence. However, in NSCLC cell lines lacking p53 or containing mutated forms of p53, compound **11** showed no substantial effects. This suggests that the quinoline-based compounds examined in our study hold potential for further development into advanced p53-targeting therapeutics for cancers harboring wild-type p53. Conducting comparative studies on different p53 mutational subtypes could enable us to tailor therapeutic strategies to each specific subtype effectively.

## Material and Methods

### Chemical synthesis

The detailed synthetic procedures used to prepare compounds **3**-**17** followed the general procedure established previously (Valente et al., 2014) and will be described in a separate medicinal chemistry paper. Other inhibitors used in this study include GSK3484862 (MedChemExpress, HY-135146), GSK3685032 (MedChemExpress, HY-139664), and GSK3735967 (MedChemExpress, HY-150249), SGI-1027 (**1**) (MedChemExpress, HY-13962), MC3343 (**2**) (ProbeChem, PC-35290). MC4741 [(2R,3S,4R,5R)-2-(Hydroxymethyl)-5-(6-((3-phenylpropyl)-amino)-9H-purin-9-yl)tetrahydrofuran-3,4-diol] is compound 14 as recently described (Zhou et al., 2023).

The DNA oligonucleotides used for substrates were synthesized by Integrated DNA Technologies (IDT) and listed in Supplementary Table S2.

### Inhibition assays of DNA methylation

The methyltransferases used in the current study were characterized previously and prepared in our laboratories: human DNMT1 residues 350-1600 (pXC915) (Hashimoto et al., 2012b), Dnmt3a2-3L (pXC465 and pXC391) and Dnmt3b2-3L (pXC273 and pXC391) (Zeng et al., 2020), *Clostridioides difficile* CamA (pXC2184) (Zhou et al., 2021), *Caulobacter crescentus* [now called *C. vibrioides*] CcrM (pXC2121) (Horton et al., 2019), and *Escherichia coli* Dam (pXC1612) (Horton et al., 2006; Horton et al., 2005; Horton et al., 2015).

The inhibition of DNA methylation was quantified using the Promega bioluminescence assays (MTase-Glo^TM^) (Hsiao et al., 2016). This assay utilizes a coupled reaction mechanism, where SAH produced during the methylation process is transformed into ATP in two distinct steps. The resultant ATP is then detected via a luciferase reaction. Luminescence signals were measured using a Synergy 4 multimode microplate reader (BioTek). Concentrations of SAH were determined based on a standard SAH curve, employing a linear regression analysis of the luminescence data. The specific reaction conditions applied for each MTase in the presence of a 10 µM concentration of the inhibitor are detailed in Supplementary Table S2.

### Inhibition assays of BER glycosylases

Generally, double-stranded DNA molecules (40 nM) featuring a single mismatch and FAM-labeled on the strand containing the mispaired nucleotide were synthesized by Integrated DNA Technologies (IDT). These were incubated with various inhibitor compounds at specified concentrations at room temperature for 10 minutes in a reaction buffer composed of 20 mM Tris (pH 8.0), 1 mM ethylenediaminetetraacetic acid (EDTA), 1 mM Tris (2-carboxyethyl) phosphine (TCEP), and 0.1 mg/ml bovine serum albumin (BSA). The reaction was initiated by adding DNA glycosylases (MBD4, TDG, or MYH) at a concentration of 400 nM. The mixtures were then incubated at room temperature for 60 minutes, and the reactions were halted by the addition of 0.1 M NaOH and subsequent heating at 95°C for 10 minutes (Haldar et al., 2022). Following this, samples were mixed with 2× loading buffer (containing 98% formamide, 1 mM EDTA, and trace amounts of bromophenol blue and xylene cyanole), heated again at 95°C for 10 minutes, and then cooled on ice. A 5-µl aliquot of each sample was loaded onto a 10 cm × 10 cm denaturing polyacrylamide gel electrophoresis (PAGE) gel, which contained 15% acrylamide, 7 M urea, and 24% formamide in 1× TBE. Electrophoresis was performed using 1X TBE buffer at 200 V for 35 minutes. Gels were scanned for visualization using a BIO-RAD ChemiDoc MP Imaging system.

For assessing UDG activity on uracil-containing DNA, both double-stranded and single-stranded, 40 nM FAM-labeled DNA was incubated at room temperature for 10 minutes, with or without 2 µM of compound **13**. The reaction was commenced by the addition of 20 nM UDG. These mixtures were then incubated at room temperature for the duration specified (from 15 sec to 60 min) and were terminated by the addition of 0.1 M NaOH and heating at 95°C for 10 minutes. The remaining steps were the same.

### Inhibition assays of Polymerases Pol η and Pol θ

Pol η and Pol θ biochemical assays, for nucleotide incorporation activity, were performed as previously described (Li et al., 2023; Wilson et al., 2021). The assays were executed using the DNA template and 5’-fluorescein-labeled DNA primer (Supplementary Table S2). Reactions were conducted at 37°C for 5 min and were stopped by adding formamide quench buffer to the final concentrations of 40% formamide, 50 mM EDTA (pH 8.0), 0.1 mg/ml xylene cyanol, and 0.1 mg/ml bromophenol. After heating to 97°C for 5 min and immediately placing on ice, reaction products were resolved on 22.5% polyacrylamide urea gels. The gels were visualized by a Sapphire Biomolecular Imager and quantified using the built-in software. Visual representation of the acquired data was rendered in Graph Prism.

The initial inhibitor screens (Figure 5A) were performed with 1 nM Pol η or Pol θ and 10 or 50 µM inhibitor, in the reaction buffer of 0.2 µM DNA, 50 µM dNTP, 150 mM KCl, 45 mM Tris (pH 7.5), 5 mM MgCl_2_, 10 mM DTT, 0.1 mg/mL bovine serum albumin, 5% glycerol, and 10% DMSO. For IC_50_ measurement (Figure 5B, C), compounds **9** and **13** were serially diluted with DMSO and added to a reaction mixture to a final concentration of 0.01-20 mM, in the reaction buffer contained 0.3-15 nM Pol η or Pol θ. For steady-state kinetics (Figure 5D-5F), the reaction mixture contained 0.2-1 nM Pol η, 120-700 µM dATP, with or without inhibitor (15 µM compound **9** or 8 µM compound **13**).

### HIV reverse transcriptase **(RT)**

HIV RT biochemical assays testing nucleotide incorporation activity were performed as previously described (Tian et al., 2018). The inhibition mixture contained 10 or 50 µM inhibitor, 10 nM HIV RT, 0.5 µM DNA, 50 µM dNTP, 50 mM KCl, 45 mM Tris (pH 7.5), 5 mM MgCl_2_, 1.3 mM DTT, 0.1 mg/mL bovine serum albumin, 4% glycerol, and 10% DMSO. The synthesis assays were executed using the RNA template and 5’-fluorescein-labeled DNA primer (Supplementary Table S2). Reactions were conducted at room temperature for 5 min and were stopped by adding formamide quench buffer to the final concentrations of 40% formamide, 50 mM EDTA (pH 8.0), 0.1 mg/ml xylene cyanol, and 0.1 mg/ml bromophenol. The gel products were analyzed similar as for Pol η.

### RNA-dependent RNA polymerase **(RdRp)**

Biochemical assays of RdRp from poliovirus testing nucleotide incorporation activity were performed as previously described (Arnold and Cameron, 2000). Each compound was assayed at two different concentrations, 10 µM and 50 µM, with 5 µM of RdRp and 1 µM of a 5’-fluorescein-labeled symmetrical self-annealing RNA template (Supplementary Table S2). The reaction mixture contained 50 mM Tris-HCl (pH 7.5), 10 mM 2-mercaptoethanol, 5 mM MgCl_2_, 60 µM ZnCl_2_, 250 µM ribonucleoside triphosphates (rNTPs), 1.3 mM DTT, 0.1 mg/mL bovine serum albumin, 4% glycerol, and 10% DMSO. A 5’-fluorescein-labeled symmetrical self-annealing RNA template (5’-GCA UGG GCC C-3’) was used. Reactions were conducted at 30 °C for 5 min and were stopped by adding formamide quench buffer to the final concentrations of 40% formamide, 50 mM EDTA (pH 8.0), 0.1 mg/ml xylene cyanol, and 0.1 mg/ml bromophenol. The gel products were analyzed similar as for Pol η.

### Isothermal titration calorimetry

The ITC experiment was performed at a constant temperature of 25 °C using the MicroCal PEAQ-ITC automated system (Malvern Instrument Ltd). Data analysis was conducted using the ITC data analysis module supplied by the manufacturer.

Double-stranded T:G mismatch oligonucleotides and compound **13** were prepared in a uniform buffer solution consisting of 20 mM Tris (pH 7.5), 150 mM NaCl, 5% glycerol, and 1% DMSO. In the sample cell, DNA was maintained at a concentration of 20 µM, and compound **13**, at a concentration of 200 µM, was injected via syringe. The process entailed thirteen injections, each delivering 3 µl of the compound into the cell. This was done with continuous stirring at a rate of 750 rpm, and the reference power was set to 8 μcal/s. Each injection lasted for 4 seconds, and there was a 200-second interval between injections to allow the system to reach equilibrium.

For interactions with a 19-bp dsDNA or CamA protein, 200 µM of compound **9**, **11**, or **13** was titrated against 20 µM of DNA or CamA. To assess the impact of compound **9** on the CamA-DNA interaction, 200 µM of compound **9** was pre-incubated separately with 200 µM of DNA or 20 µM of CamA in a buffer containing 150 mM NaCl, 20 mM Tris-HCl pH 7.5, 0.5 mM TCEP, and 1% DMSO.

### Southern blot DNA methylation assays

Analysis of DNA methylation at the minor satellite repeats in mouse embryonic stem cells was carried out as described previously (Veland et al., 2019). Cells were cultured in gelatin-coated petri dishes in DMEM supplemented with 15% FBS, 0.1 mM nonessential amino acids, 0.1 mM ß-mercaptoethanol, 1% penicillin/streptomycin, and 10^3^ U/ml leukemia inhibitory factor (LIF). Cells were seeded onto 6-well plates at a cell density of ∼5 X 10^5^/well, and cells were treated with 0.1% DMSO (control) or at 0.1 µM for GSK3685032 and GSK3484862, and 2 µM for quinoline-based compounds for 48 h. Genomic DNA (1 µg) was digested with the methylation-sensitive restriction enzyme HpaII (New England Biolabs) and analyzed by Southern hybridization with a specific biotin-labeled DNA probe (300 ng) (Supplementary Table S2). Detection was performed using the North2South Chemiluminescent Hybridization and Detection Kit (Thermo Fisher Scientific).

For analysis of whole cell extracts by Western blot, mESCs were lysed in cold RIPA buffer [50 mM Tris–HCl (pH 8.8), 150 mM NaCl, 1% Triton X-100, 0.5% Sodium Deoxycholate, 0.1% SDS, 1 mM EDTA, 3 mM MgCl2, and 1 X protease inhibitor cocktail (Thermo Fisher Scientific)]. The antibodies used were DNMT1 (Cell Signaling Technology (CST), #5032) and mouse p53 (NCL-L-p53-CM5p, Leica Biosystems).

### X-ray crystallography

For crystallization, the CamA-DNA-inhibitor complexes were prepared following previously established methods (Zhou et al., 2021; Zhou et al., 2023). In brief, crystal growth was achieved under similar conditions: 21−24% (w/v) polyethylene glycol 3350, 0.1 M Tris-HCl with a pH range of 7.0−7.5, and 0.28 M potassium citrate, all at room temperature (∼19 °C). The crystals typically appeared after 3−4 days, using the sitting drop vapor diffusion technique. An Art Robbins Gryphon Crystallization Robot was employed for setting up 0.4 µL drops. Notably, crystals formed with inhibitors 455 or 462 at a concentration of 200 µM exhibited a distinct shape compared to those without inhibitors. For preservation, these crystals were rapidly frozen in liquid nitrogen following a brief immersion in the reservoir solution, which had been fortified with 20% (v/v) ethylene glycol.

Diffraction data for the crystals were collected at the SER-CAT beamline 22ID of the Advanced Photon Source at Argonne National Laboratory. The crystals were maintained in a cryostream at 100 K, and data collection typically involved rotating each crystal by 0.5° for each of the 800 frames. The crystallographic datasets were processed using HKL2000 (Otwinowski et al., 2003) (Supplementary Table S1).

Structures of the CamA-DNA-inhibitor ternary complex were resolved using the difference Fourier method (Liebschner et al., 2017). For crystals incorporating only compounds **9** or **11**, our previously determined binary CamA-DNA structure (PDB ID: 7LNJ) was used as a search model. For the crystal containing both compounds **9** and MC4741, the CamA-DNA-MC4741 structure (PDB ID: 8CXZ) served as the starting model. Given that the unit cell parameters of all crystals were essentially isomorphous to those of previously obtained crystals, rigid body refinement was employed in the initial refinement cycle to position the new structures within the unit cell. Difference electron density maps (2Fo-Fc and Fo-Fc) were utilized to locate the bound compounds **9** or **11**. For each compound, a SMILES string was submitted to the Grade web server (http://grade.globalphasing.org) to generate geometrical restraints, which were provided in a CIF file. This file was used in subsequent refinement cycles and also facilitated the provision of the compound’s structure in PDB format.

Initially, the extra densities in the crystal structures suggested the possibility of several conformations for the compounds. However, only the most distinct and likely predominant conformation was modeled and refined within the structure. All refinements were carried out using PHENIX REFINE (Afonine et al., 2012), which included 5% of reflections randomly selected for validation, as indicated by the R-free value (Brunger, 1997) (Supplementary Table S1). The quality of the structures was continually assessed during the PHENIX refinement process and supplemented with manual inspection using COOT (Emsley and Cowtan, 2004). The final structure models underwent validation by the PDB validation server (Read et al., 2011). Images of the structures were generated using PyMol (Schrödinger, LLC).

### Compound treatments in cell lines

Four lung cancer cell lines (A549-Luc2, NCI-H1299, NCI-H596 and NCI-H1975), osteosarcoma U2OS, breast cancer MCF7 and prostate cancer PC3 cell lines were purchased from the American Type Culture Collection (ATCC) and validated at The University of Texas MD Anderson Cancer Center (Houston, TX). A549-Luc2, U2OS and MCF7 contain the wild-type p53 gene; H1299 and PC3 are p53-null; H596 and H1975 contain a hotspot p53 mutation (p.G245C for H596, p.R273H for H1975). All cells were incubated at 37°C with 5% CO_2_. A549-Luc2 cells were cultured in ATCC-formulated F-12K Medium (Catalog No. 30-2004) supplemented with 10% fetal bovine serum (FBS) (Sigma-Aldrich) and 1% penicillin/streptomycin. MCF7, U2OS and PC3 cells were cultured in Dulbecco’s Modified Eagle’s Medium (DMEM) with L-glutamine and 4.5g glucose/L but without sodium pyruvate (Mediatech) supplemented with 10% FBS and 1% penicillin/streptomycin. NCI-H1299, NCI-H596 and NCI-H1975 cells were cultured in ATCC-formulated RPMI-1640 Medium (ATCC 30-2001) supplemented with 10% FBS and 1% penicillin/streptomycin. Cells were seeded onto 6-well plates at a cell density of ∼5×10^5^/well. The next day, cells were treated with 0.1% DMSO (control) or compounds at the indicated concentrations and duration. Cells were collected for western blot assay.

Cells were lysed with sodium dodecyl sulfate (SDS) sample buffer. The lysates were separated by Bis-Tris sodium dodecyl sulfate polyacrylamide gel electrophoresis (SDS-PAGE) using 4–20% precast polyacrylamide gel (BioRad, #4561096). The proteins were transferred to low fluorescence polyvinylidene difluoride (PVDF) membranes (BioRad, #1620261), which were blocked with 5% non-fat dry milk in Tris-buffered saline with Tween 20 (TBST) at room temperature for 1 h and then probed with primary followed by secondary antibodies. The signals were detected with Clarity Western ECL substrate (Bio-Rad Laboratories, #1705061) and imaged using a ChemiDoc imaging system (Bio-Rad Laboratories).

The primary antibodies used in this study: DNMT1 [(Cell Signaling Technology (CST), #5032), DNMT3A (CST, #3598), p53 (DO-1; Santa Cruz, sc-126), phosphor p53 (Ser15) (Abcam, ab1431), p21 (BD Biosciences, #556431), Vinculin (Sigma, SAB4200729), H3 (CST, #14269), GAPDH (CST, #2118), Actin (Sigma-Aldrich, A2228). The secondary antibodies used were: HRP-conjugated anti-rabbit-IgG (CST, #7074) and HRP-conjugated anti-mouse-IgG (Abcam, ab6820).

For cell viability assay, A549-Luc2 cells were seeded onto 96-well plates at a cell density of ∼1×10^4^/well. The following day, 0.1% DMSO or compounds at the indicated concentrations were added and incubated for 3 days. The viability was measured by using the CellTiter-Glo® Luminescent Cell Viability Assay (Promega, G7572).

## Data Availability Statement

The experimental data that support the findings of this study are contained within the article. The authors have deposited the X-ray structure (coordinates) and the source data (structure factor file) of CamA-DNA with bound inhibitors to the PDB and will be released upon article publication under accession numbers PDB 8VPG (compound **9**), PDB 8VPI (compound **11**), and PDB 8VPH (compounds **11** and MC4741).

## Supporting Information

**Supplementary Figures1 S1-S6 and Tables S1-S2.**

## Author Contributions

J.Z. performed CamA and CcrM protein purification, inhibition assays, and crystallization. Q.C. performed cellular inhibition in cancer cell lines. J.Z. and J.R.H. performed X-ray crystallography experiments. R.R. performed protein purifications of DNMTs and their inhibition assays. J.R.H. provided Dam enzyme used in the study. J.Y. performed inhibition assays of DNA glycosylases and ITC experiments. B.L. performed experiments in mESCs. C.C. performed inhibition assays of Pol eta and HIV RT. C.L. and L.M. performed inhibition assays of Pol theta. Y.Y. performed inhibition assays of RNA-dependent RNA polymerase. D.R. provided compound MC4741. R.M.B. participated in discussion and assisted in preparing the manuscript. Y.G. guided inhibition of polymerases. S.V. and A.M. designed and provided the compounds **3-17**. X.Z., T.C. and X.C. organized and designed the scope of the study.

## Notes

The authors declare no competing financial interest.

## Acknowledgements

We thank Craig E. Cameron for provide plasmid of poliovirus RNA-dependent RNA polymerase; Mateo Ramírez-Valentini for help with the pol eta assay; Abhinav Jain for advice on p53 biology and provide mouse p53 antibody. We thank the beamline scientists of Southeast Regional Collaborative Access Team (SER-CAT) at the Advanced Photon Source (APS), Argonne National Laboratory, USA. The use of SER-CAT is supported by its member institutions and equipment grants (S10_RR25528, S10_RR028976, and S10_OD027000) from the US National Institutes of Health. Use of the APS was supported by the U.S. Department of Energy, Office of Science, Office of Basic Energy Sciences, under contract W-31-109-Eng-38. The work was supported by U.S. National Institutes of Health grant R35GM134744 (to X.C.), R21CA277152 (to X.C. and T.C.), Cancer Prevention and Research Institute of Texas grant RR160029 (to X.C., who is a CPRIT Scholar in Cancer Research), a Developmental Research Program grant (A22-0002-S013) of NCI SPORE Project 5P50CA254897 (administered via Coriell Institute for Medical Research), the Cockrell Foundation in Houston (to J. Z.), and AIRC (Associazione Italiana per la Ricerca sul Cancro) (IG26172) and Ateneo Sapienza Project 2020 (RG120172B8E53D03) to S.V, and FISR2019_00374 MeDyCa to A.M.

## Abbreviations

The abbreviations used are

MTases: methyltransferases
SAM: *S*-adenosyl-L-methionine
SAH: *S*-adenosyl-L-homocysteine
5mC: 5-methylcytosine
6mA: 6-methyladenine

## Footnotes

1 https://www.fda.gov/drugs/resources-information-approved-drugs/fda-approves-azacitidine-newly-diagnosed-juvenile-myelomonocytic-leukemia

2 https://www.fda.gov/drugs/resources-information-approved-drugs/fda-approves-oral-combination-decitabine-and-cedazuridine-myelodysplastic-syndromes

---

**Figure S1.**
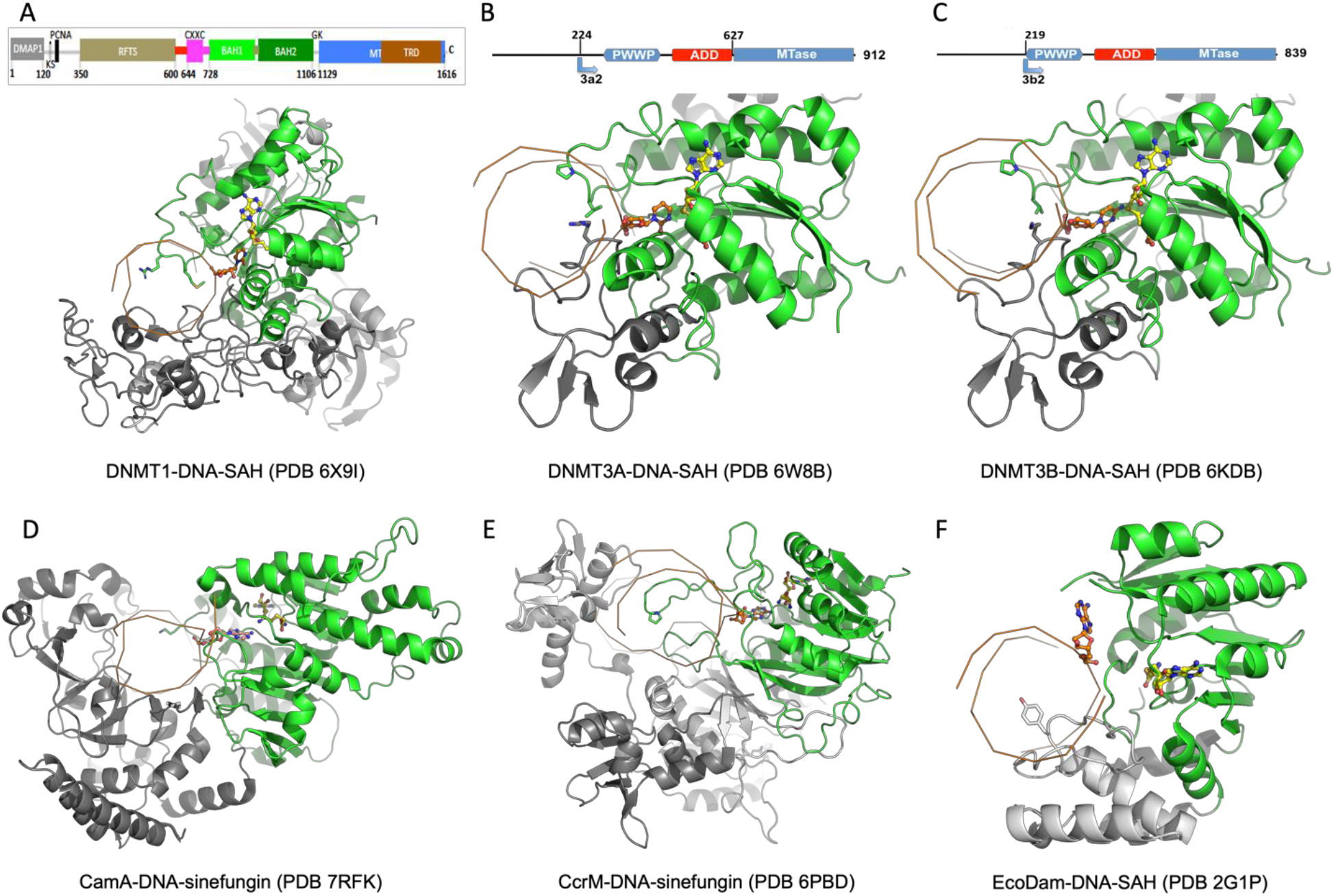
The structures of six MTases. (**A**) human DNMT1 (PDB 6X9I) with schematic domain arrangement shown above, (**B**) human DNMT3A (PDB 6W8B) with schematic domain arrangement shown above, (**C**) human DNMT3B (PDB 6KDB) with schematic domain arrangement shown above, (**D**) *C. difficile* CamA (PDB 7RFK), (**E**) *C. crescentus* CcrM (PDB 6PBD), and (**F**) *E. coli* Dam (PDB 2G1P). The conserved catalytic MTase domains are colored in green, whereas the DNA recognition domains are in grey. For simplicity, the DNA strands are shown as orange ribbons (with the helical axis projecting out of the page), and the flipped target nucleotide and SAH (or sinefungin) in the SAM binding pocket are in ball-and-stick. The protein residues that intercalate into the DNA are shown in stick models.

**Figure S2.**
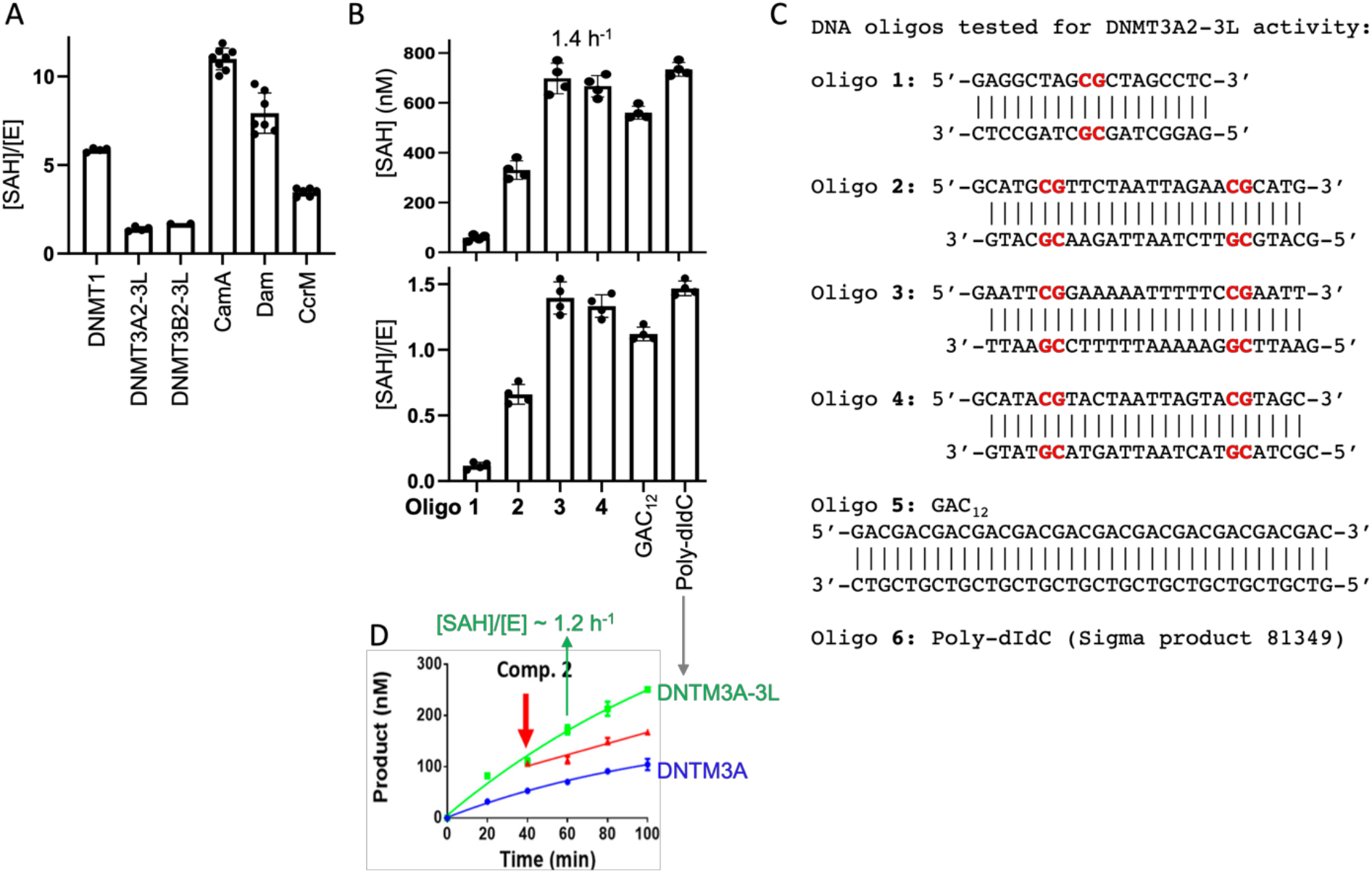
Related to Figure 1. (**A**) The enzymatic turnover number of six MTases under the conditions used (without any inhibitor), presented as the formation of byproduct SAH concentration per enzyme molecule [SAH]/[E], in a bioluminescence assay. DNMT3A2-3L and DNMT3B2-3L have low turnover numbers. (**B-C**) Comparison of DNMT3A2-3L activity on six DNA substrates. The best value of apparent *k*cat is ∼1.4 h^−1^ for oligos 3 and 4 as well as poly-dIdC. (**D**) Adapted from an earlier study (Sandoval et al., 2022). Sandoval et al. commented that “Poly dI-dC was used as a substrate due to the increased activity of DNMT3A”, which resulted in an apparent *k*_cat_ of ∼1.2 h^−1^.

**Figure S3.**
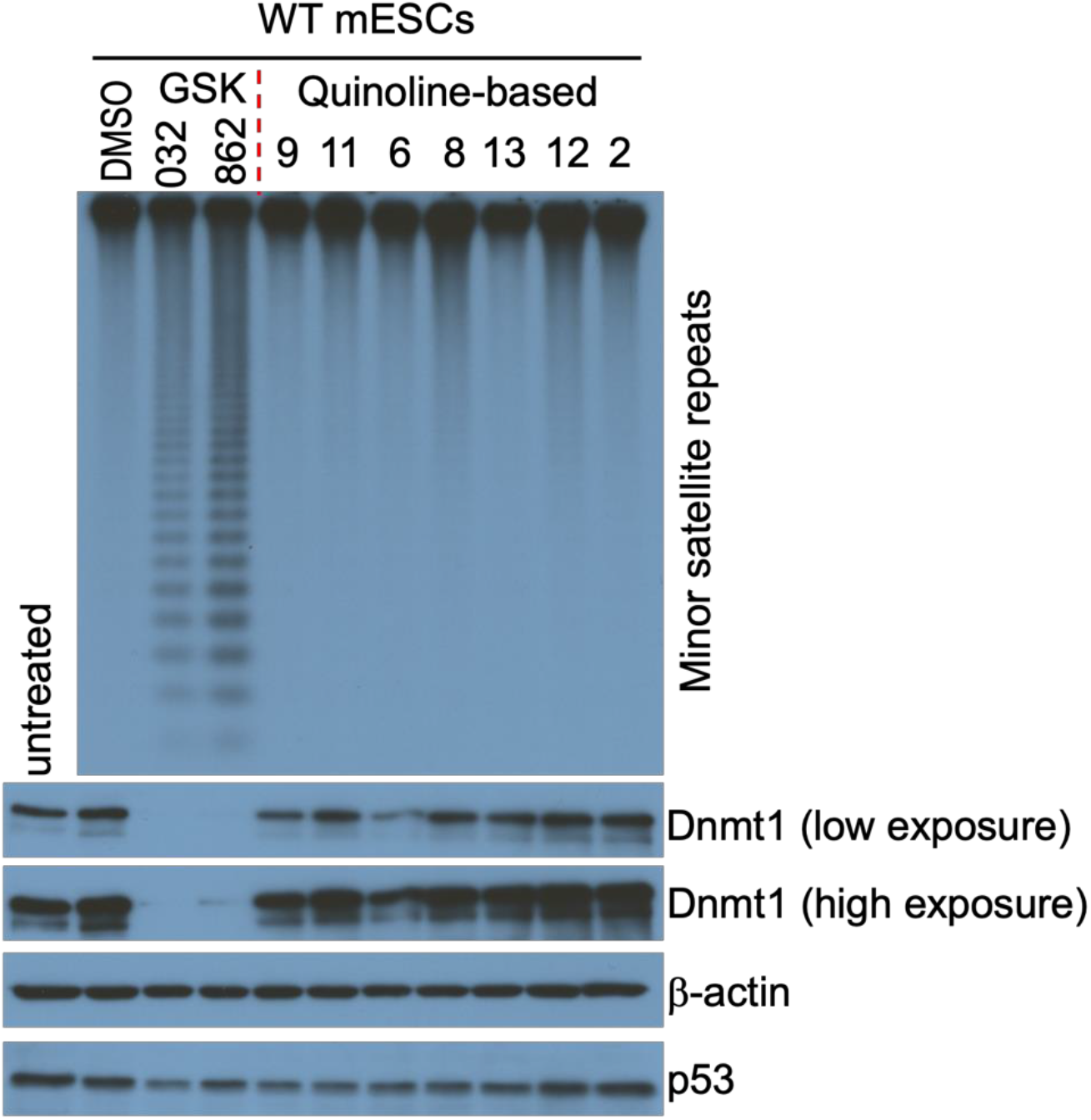
Related to Figure 2. WT mESCs were cultured with inhibitors or DMSO for two days, and genomic DNA was analyzed for methylation at the minor satellite repeats by Southern blot after digestion with the methylation-sensitive enzyme *Hpa*II. Six quinoline-based derivatives at 2 μM and two GSK DNMT1 inhibitors (GSK3685<u>032</u> and GSK3484<u>862</u>) at 0.1 μM are compared. We note that treatment with compound **6** decreases DNMT1 level significantly.

**Figure S4.**
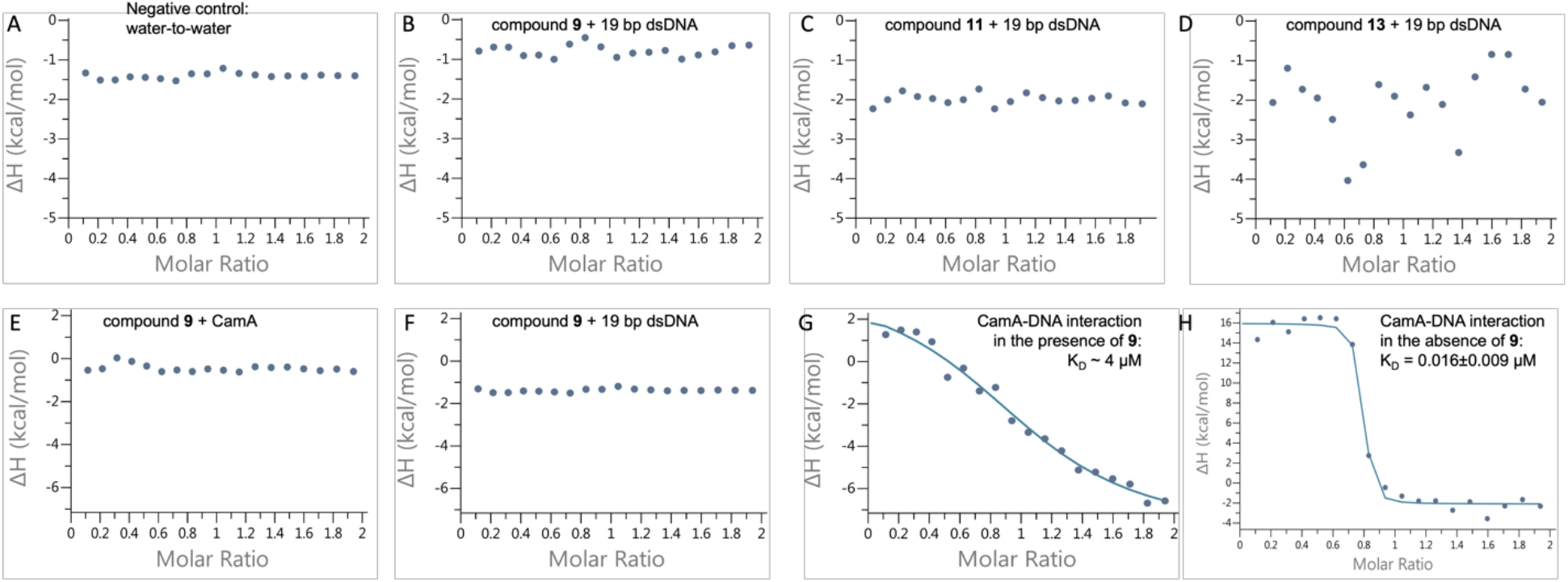
Related to Figure 3. No direct interaction between 19-bp DNA oligonucleotide and the quinoline compounds by isothermal titration calorimetry (ITC) measurements. (**A**) Water-to-water control. (**B**-**D**) No interaction between compounds **9**, **11** or **13** and DNA. (**E**) No interaction between compound **9** and CamA. (**F**) No interaction between compound **9** and DNA. (**G**) CamA and DNA have weak interactions in the presence of compound **9** (K_D_ ∼ 4 μM). (**H**) CamA and DNA have strong interactions in the absence of compound **9** (K_D_=0.16 μM).

**Figure S5.**
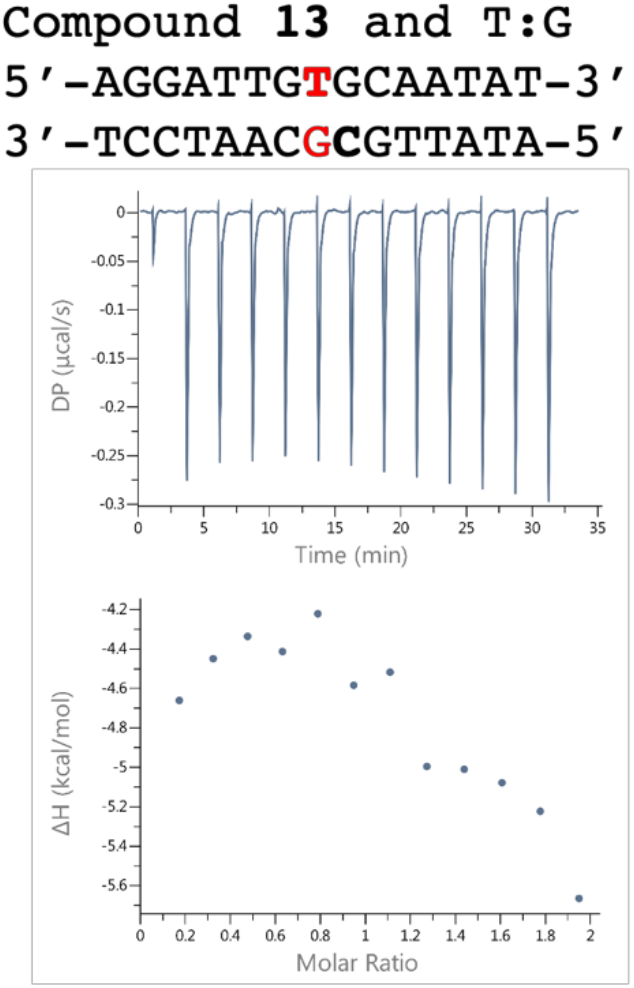
Related to Figure 4. Isothermal titration calorimetry (ITC) measurement showed no direct interaction between compound **13** and dsDNA containing a single T:G mismatch.

**Figure S6.**
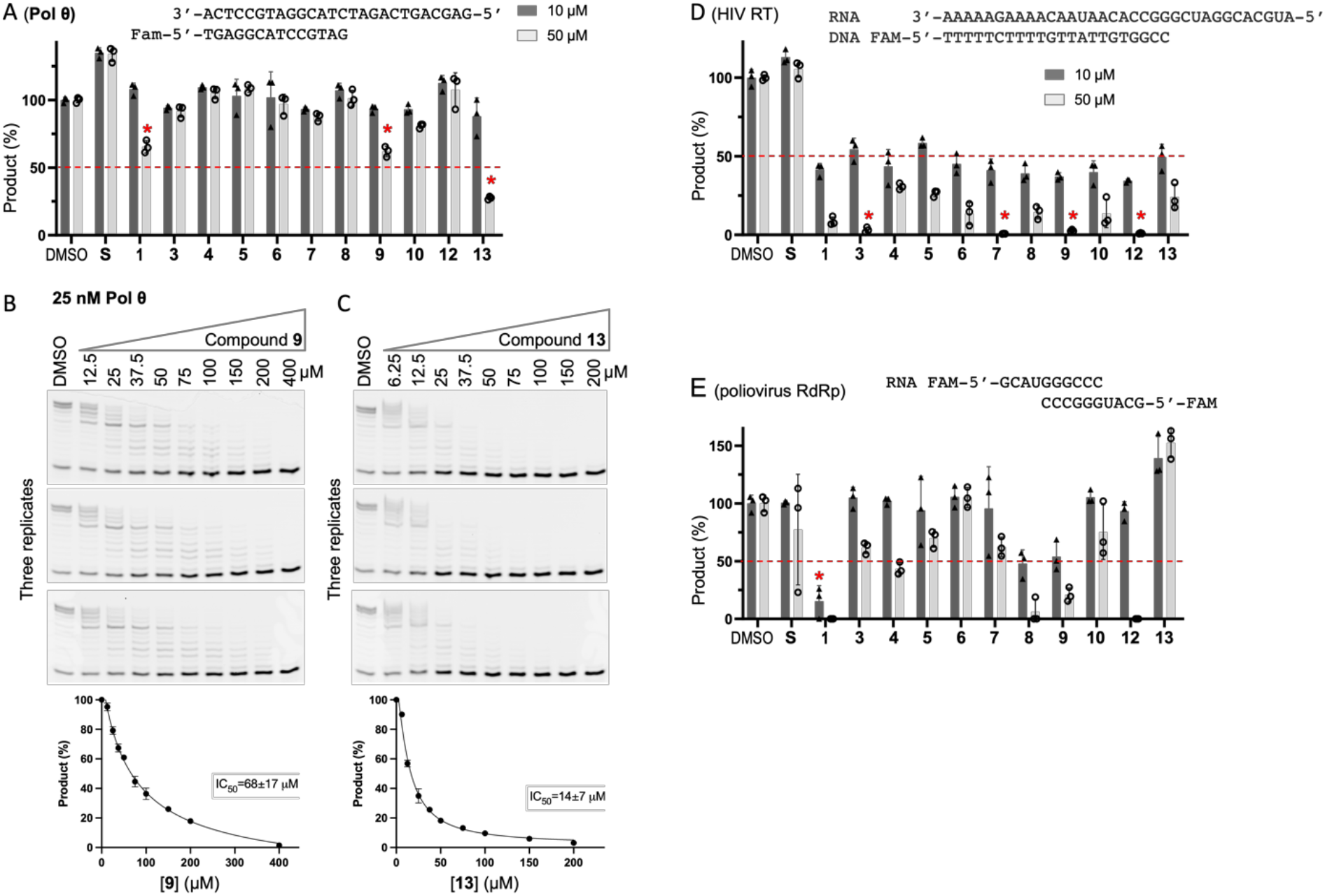
Related to Figure 5. Inhibition of other polymerases. (**A**) Relative inhibition of Pol θ at two inhibitor concentrations of 10 and 50 μM. [S, sinefungin; compounds **2** and **11** were unavailable at the time of these assays.] (**B-C**) Measurement of IC_50_ values of compound **9** (panel B) and compound **13** (panel C). Three replicates of reactions with increased compound concentrations, reaction products were resolved on 22.5% polyacrylamide urea gels. The gels were visualized and quantified (bottom panels). (**D-E**) Inhibition screen of the quinoline-based compounds at two concentrations (10 µM and 50 µM) against HIV reverse transcriptase (RT) (panel D), and poliovirus RNA-dependent RNA polymerase (RdRp) (panel E).

**Table S1.** Summary of X-ray data collection at wavelength=1Å and refinement statistics (*)

|  |  |  |  |
| --- | --- | --- | --- |
| Beamline | APS 22-ID |  |  |
| Date collected | 09/19/2021 | 11/03/2021 | 09/19/2021 |
| Inhibitor | 455 | 455 + 174 | 462 |
| PDB code | 8VPG | 8VPH | 8VPI |
| Space group | $P2_12_12_1$ | $P2_12_12_1$ | $P2_12_12_1$ |
| Cell dimensions (Å) | 80.54, 161.60, 234.98 | 81.59, 161.71, 235.45 | 81.59, 161.26, 235.72 |
| Resolution (Å) | 41.84-3.05<br>(3.16-3.05) | 45.78-3.18<br>(3.30-3.18) | 47.77-2.89<br>(2.99-2.89) |
| <sup>a</sup> R<sub>merge</sub> | 0.384 (3.36) | 0.413 (2.86) | 0.321 (2.66) |
| R<sub>pim</sub> | 0.085 (0.917) | 0.118 (0.999) | 0.093 (1.01) |
| CC<sub>1/2</sub>, CC* | (0.394, 0.752) | (0.357, 0.725) | (0.309, 0.687) |
| <sup>b</sup> <I/σI> | 10.2 (0.9) | 8.2 (1.0) | 8.5 (0.7) |
| Completeness (%) | 99.9 (99.5) | 99.9 (100.0) | 98.5 (88.4) |
| Redundancy | 20.8 (13.3) | 13.2 (8.5) | 11.3 (6.1) |
| Observed reflections | 1,230,186 | 696,323 | 782,701 |
| Unique reflections | 59,233 | 52,804 | 69,180 |
| **Refinement** |  |  |  |
| Resolution (Å) | 3.05 | 3.18 | 2.89 |
| No. reflections | 59,012 | 52,693 | 68,885 |
| <sup>c</sup> R<sub>work</sub> / <sup>d</sup> R<sub>free</sub> | 0.196 / 0.231 | 0.176 / 0.231 | 0.196 / 0.241 |
| No. Atoms |  |  |  |
| Protein | 13,334 | 13,332 | 13,324 |
| DNA | 1,686 | 1,686 | 1,686 |
| Inhibitor | 39 | 123 | 43 |
| Solvent | 65 | 55 | 66 |
| B Factors (Å<sup>2</sup>) |  |  |  |
| Protein | 96.6 | 91.4 | 90.2 |
| DNA | 109.5 | 106.2 | 103.4 |
| Inhibitor | 132.8 | 110.6 | 120.4 |
| Solvent | 64.8 | 58.5 | 64.6 |
| **R.m.s. deviations** |  |  |  |
| Bond lengths (Å) | 0.003 | 0.004 | 0.002 |
| Bond angles (°) | 0.5 | 0.6 | 0.5 |

\* Values in parenthesis correspond to highest resolution shell.
<sup>a</sup> R<sub>merge</sub> = $\sum |I - \langle I \rangle| / \sum I$ , where I is the observed intensity and <I> is the averaged intensity from multiple observations.
<sup>b</sup> <I/σI> = averaged ratio of the intensity (I) to the error of the intensity (σI).
<sup>c</sup> R<sub>work</sub> = $\sum |F_{obs} - F_{cal}| / \sum |F_{obs}|$ , where F<sub>obs</sub> and F<sub>cal</sub> are the observed and calculated structure factors, respectively.
<sup>d</sup> R<sub>free</sub> was calculated using a randomly chosen subset (5%) of the reflections not used in refinement.

**Table S2.**
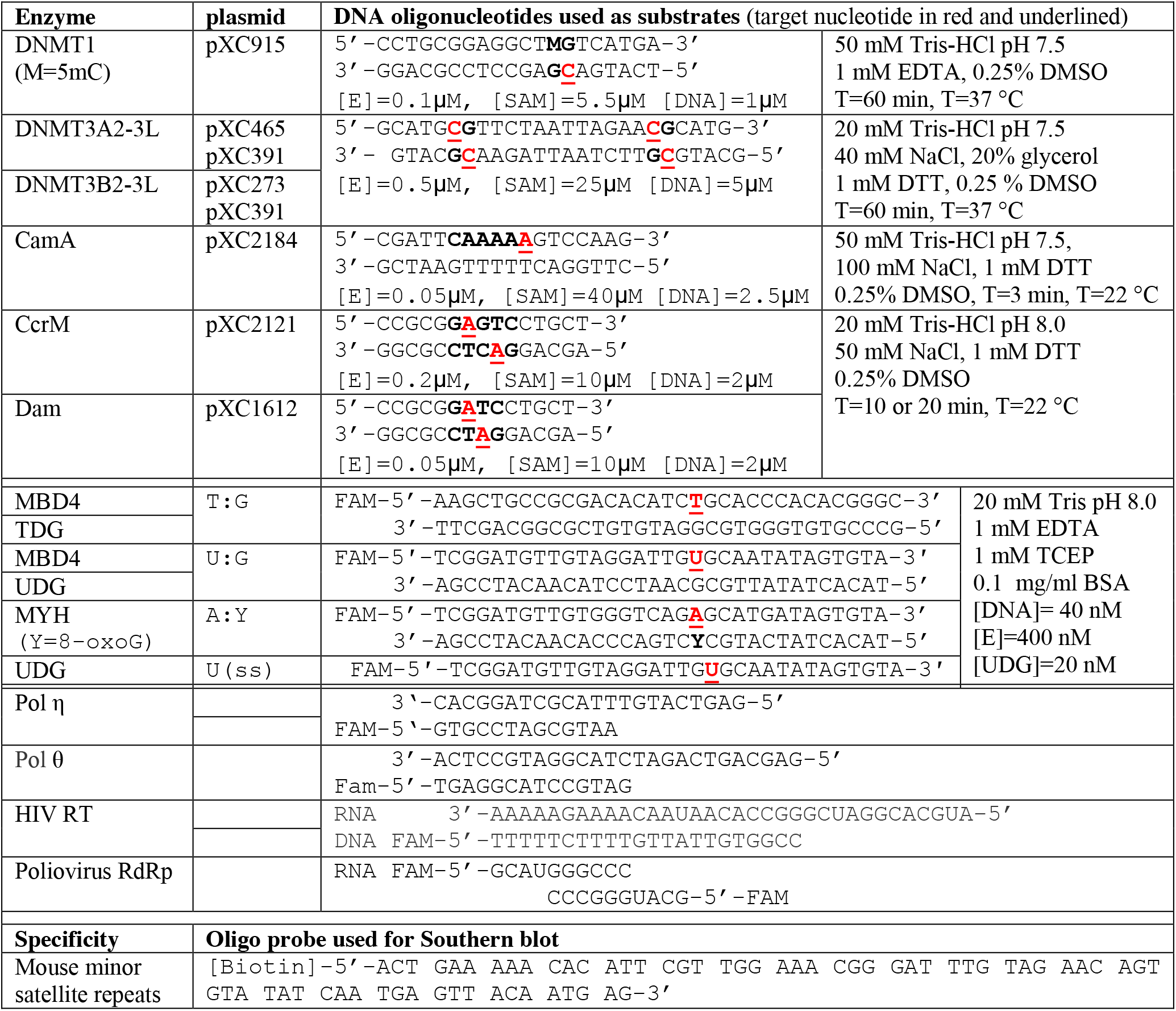
DNA oligonucleotides and reaction conditions used in this study.

## References

Abuetabh, Y., Wu, H.H., Chai, C., Al Yousef, H., Persad, S., Sergi, C.M., and Leng, R. (2022). DNA damage response revisited: the p53 family and its regulators provide endless cancer therapy opportunities. Exp Mol Med 54, 1658–1669.

Afonine, P.V., Grosse-Kunstleve, R.W., Echols, N., Headd, J.J., Moriarty, N.W., Mustyakimov, M., Terwilliger, T.C., Urzhumtsev, A., Zwart, P.H., and Adams, P.D. (2012). Towards automated crystallographic structure refinement with phenix.refine. Acta Crystallogr D Biol Crystallogr 68, 352–367.

Allis, C.D., Jenuwein, T., and Reinberg, D. (2015). Epigenetics, second edition (Cold Spring Harbor, NY: Cold Spring Harbor Laboratory Press).

Arnold, J.J., and Cameron, C.E. (2000). Poliovirus RNA-dependent RNA polymerase (3D(pol)). Assembly of stable, elongation-competent complexes by using a symmetrical primer-template substrate (sym/sub). J Biol Chem 275, 5329–5336.

Ayaz, G., Yan, H., Malik, N., and Huang, J. (2022). An Updated View of the Roles of p53 in Embryonic Stem Cells. Stem Cells 40, 883–891.

Besselink, N., Keijer, J., Vermeulen, C., Boymans, S., de Ridder, J., van Hoeck, A., Cuppen, E., and Kuijk, E. (2023). The genome-wide mutational consequences of DNA hypomethylation. Scientific reports 13, 6874.

Briski, R., Garcia-Manero, G., Kantarjian, H., and Ravandi, F. (2023). The history of oral decitabine/cedazuridine and its potential role in acute myeloid leukemia. Ther Adv Hematol 14, 20406207231205429.

Brunger, A.T. (1997). Free R value: cross-validation in crystallography. Methods Enzymol 277, 366–396.

Carroll, A.G., Voeller, H.J., Sugars, L., and Gelmann, E.P. (1993). p53 oncogene mutations in three human prostate cancer cell lines. The Prostate 23, 123–134.

Chen, Q., Liu, B., Zeng, Y., Hwang, J.W., Dai, N., Correa, I.R., Jr., Estecio, M.R., Zhang, X., Santos, M.A., Chen, T., et al. (2023). GSK-3484862 targets DNMT1 for degradation in cells. NAR Cancer 5, zcad022.

Chen, T., Ueda, Y., Dodge, J.E., Wang, Z., and Li, E. (2003). Establishment and maintenance of genomic methylation patterns in mouse embryonic stem cells by Dnmt3a and Dnmt3b. Mol Cell Biol 23, 5594–5605.

Chen, T., Ueda, Y., Xie, S., and Li, E. (2002). A novel Dnmt3a isoform produced from an alternative promoter localizes to euchromatin and its expression correlates with active de novo methylation. J Biol Chem 277, 38746–38754.

Cheng, X., and Blumenthal, R.M. (2002). Cytosines do it, thymines do it, even pseudouridines do it--base flipping by an enzyme that acts on RNA. Structure 10, 127–129.

Cristalli, C., Manara, M.C., Valente, S., Pellegrini, E., Bavelloni, A., De Feo, A., Blalock, W., Di Bello, E., Pineyro, D., Merkel, A., et al. (2022). Novel Targeting of DNA Methyltransferase Activity Inhibits Ewing Sarcoma Cell Proliferation and Enhances Tumor Cell Sensitivity to DNA Damaging Drugs by Activating the DNA Damage Response. Front Endocrinol (Lausanne) 13, 876602.

Datta, J., Ghoshal, K., Denny, W.A., Gamage, S.A., Brooke, D.G., Phiasivongsa, P., Redkar, S., and Jacob, S.T. (2009). A new class of quinoline-based DNA hypomethylating agents reactivates tumor suppressor genes by blocking DNA methyltransferase 1 activity and inducing its degradation. Cancer Res 69, 4277–4285.

Dong, G., Yasgar, A., Peterson, D.L., Zakharov, A., Talley, D., Cheng, K.C., Jadhav, A., Simeonov, A., and Huang, R. (2020). Optimization of High-Throughput Methyltransferase Assays for the Discovery of Small Molecule Inhibitors. ACS Comb Sci 22, 422–432.

Emsley, P., and Cowtan, K. (2004). Coot: model-building tools for molecular graphics. Acta Crystallogr D Biol Crystallogr 60, 2126–2132.

Esteller, M. (2007). Epigenetic gene silencing in cancer: the DNA hypermethylome. Hum Mol Genet 16 Spec No 1, R50–59.

Fenaux, P., Mufti, G.J., Hellstrom-Lindberg, E., Santini, V., Finelli, C., Giagounidis, A., Schoch, R., Gattermann, N., Sanz, G., List, A., et al. (2009). Efficacy of azacitidine compared with that of conventional care regimens in the treatment of higher-risk myelodysplastic syndromes: a randomised, open-label, phase III study. Lancet Oncol 10, 223–232.

Flotho, C., Claus, R., Batz, C., Schneider, M., Sandrock, I., Ihde, S., Plass, C., Niemeyer, C.M., and Lubbert, M. (2009). The DNA methyltransferase inhibitors azacitidine, decitabine and zebularine exert differential effects on cancer gene expression in acute myeloid leukemia cells. Leukemia 23, 1019–1028.

Gonzalez-Zulueta, M., Bender, C.M., Yang, A.S., Nguyen, T., Beart, R.W., Van Tornout, J.M., and Jones, P.A. (1995). Methylation of the 5’ CpG island of the p16/CDKN2 tumor suppressor gene in normal and transformed human tissues correlates with gene silencing. Cancer Res 55, 4531–4535.

Greger, V., Passarge, E., Hopping, W., Messmer, E., and Horsthemke, B. (1989). Epigenetic changes may contribute to the formation and spontaneous regression of retinoblastoma. Hum Genet 83, 155–158.

Gros, C., Fleury, L., Nahoum, V., Faux, C., Valente, S., Labella, D., Cantagrel, F., Rilova, E., Bouhlel, M.A., David-Cordonnier, M.H., et al. (2015). New insights on the mechanism of quinoline-based DNA Methyltransferase inhibitors. J Biol Chem 290, 6293–6302.

Gujar, H., Weisenberger, D.J., and Liang, G. (2019). The Roles of Human DNA Methyltransferases and Their Isoforms in Shaping the Epigenome. Genes (Basel) 10.

Haldar, T., Jha, J.S., Yang, Z., Nel, C., Housh, K., Cassidy, O.J., and Gates, K.S. (2022). Unexpected Complexity in the Products Arising from NaOH-, Heat-, Amine-, and Glycosylase-Induced Strand Cleavage at an Abasic Site in DNA. Chemical research in toxicology 35, 218–232.

Hashimoto, H., Hong, S., Bhagwat, A.S., Zhang, X., and Cheng, X. (2012a). Excision of 5-hydroxymethyluracil and 5-carboxylcytosine by the thymine DNA glycosylase domain: its structural basis and implications for active DNA demethylation. Nucleic Acids Res 40, 10203–10214.

Hashimoto, H., Liu, Y., Upadhyay, A.K., Chang, Y., Howerton, S.B., Vertino, P.M., Zhang, X., and Cheng, X. (2012b). Recognition and potential mechanisms for replication and erasure of cytosine hydroxymethylation. Nucleic Acids Res 40, 4841–4849.

Hashimoto, H., Zhang, X., and Cheng, X. (2012c). Excision of thymine and 5-hydroxymethyluracil by the MBD4 DNA glycosylase domain: structural basis and implications for active DNA demethylation. Nucleic Acids Res 40, 8276–8284.

Herman, J.G., Latif, F., Weng, Y., Lerman, M.I., Zbar, B., Liu, S., Samid, D., Duan, D.S., Gnarra, J.R., and Linehan, W.M. (1994). Silencing of the VHL tumor-suppressor gene by DNA methylation in renal carcinoma. Proc Natl Acad Sci U S A 91, 9700–9704.

Herman, J.G., Merlo, A., Mao, L., Lapidus, R.G., Issa, J.P., Davidson, N.E., Sidransky, D., and Baylin, S.B. (1995). Inactivation of the CDKN2/p16/MTS1 gene is frequently associated with aberrant DNA methylation in all common human cancers. Cancer Res 55, 4525–4530.

Holliday, R. (1996). DNA methylation in eukaryotes: 20 years on. Epigenetic Mechanisms of gene regulation, 5–27.

Hong, S., and Cheng, X. (2016). DNA Base Flipping: A General Mechanism for Writing, Reading, and Erasing DNA Modifications. Advances in experimental medicine and biology 945, 321–341.

Horton, J.R., Liebert, K., Bekes, M., Jeltsch, A., and Cheng, X. (2006). Structure and substrate recognition of the Escherichia coli DNA adenine methyltransferase. J Mol Biol 358, 559–570.

Horton, J.R., Liebert, K., Hattman, S., Jeltsch, A., and Cheng, X. (2005). Transition from nonspecific to specific DNA interactions along the substrate-recognition pathway of dam methyltransferase. Cell 121, 349–361.

Horton, J.R., Pathuri, S., Wong, K., Ren, R., Rueda, L., Fosbenner, D.T., Heerding, D.A., McCabe, M.T., Pappalardi, M.B., Zhang, X., et al. (2022). Structural characterization of dicyanopyridine containing DNMT1-selective, non-nucleoside inhibitors. Structure 30, 793–802.

Horton, J.R., Woodcock, C.B., Opot, S.B., Reich, N.O., Zhang, X., and Cheng, X. (2019). The cell cycle-regulated DNA adenine methyltransferase CcrM opens a bubble at its DNA recognition site. Nat Commun 10, 4600.

Horton, J.R., Zhang, X., Blumenthal, R.M., and Cheng, X. (2015). Structures of Escherichia coli DNA adenine methyltransferase (Dam) in complex with a non-GATC sequence: potential implications for methylation-independent transcriptional repression. Nucleic Acids Res 43, 4296–4308.

Hsiao, K., Zegzouti, H., and Goueli, S.A. (2016). Methyltransferase-Glo: a universal, bioluminescent and homogenous assay for monitoring all classes of methyltransferases. Epigenomics 8, 321–339.

Isaacs, W.B., Carter, B.S., and Ewing, C.M. (1991). Wild-type p53 suppresses growth of human prostate cancer cells containing mutant p53 alleles. Cancer Res 51, 4716–4720.

Jamshidi, A., Liu, M.C., Klein, E.A., Venn, O., Hubbell, E., Beausang, J.F., Gross, S., Melton, C., Fields, A.P., Liu, Q., et al. (2022). Evaluation of cell-free DNA approaches for multi-cancer early detection. Cancer Cell 40, 1537–1549 e1512.

Jia, D., Jurkowska, R.Z., Zhang, X., Jeltsch, A., and Cheng, X. (2007). Structure of Dnmt3a bound to Dnmt3L suggests a model for de novo DNA methylation. Nature 449, 248–251.

Juttermann, R., Li, E., and Jaenisch, R. (1994). Toxicity of 5-aza-2’-deoxycytidine to mammalian cells is mediated primarily by covalent trapping of DNA methyltransferase rather than DNA demethylation. Proc Natl Acad Sci U S A 91, 11797–11801.

Lange, S.S., Takata, K., and Wood, R.D. (2011). DNA polymerases and cancer. Nat Rev Cancer 11, 96–110.

Li, C., Zhu, H., Jin, S., Maksoud, L.M., Jain, N., Sun, J., and Gao, Y. (2023). Structural basis of DNA polymerase theta mediated DNA end joining. Nucleic Acids Res 51, 463–474.

Li, E., Bestor, T.H., and Jaenisch, R. (1992). Targeted mutation of the DNA methyltransferase gene results in embryonic lethality. Cell 69, 915–926.

Liebschner, D., Afonine, P.V., Moriarty, N.W., Poon, B.K., Sobolev, O.V., Terwilliger, T.C., and Adams, P.D. (2017). Polder maps: improving OMIT maps by excluding bulk solvent. Acta Crystallogr D Struct Biol 73, 148–157.

Lobner-Olesen, A., Skovgaard, O., and Marinus, M.G. (2005). Dam methylation: coordinating cellular processes. Current opinion in microbiology 8, 154–160.

Lubbert, M., Suciu, S., Baila, L., Ruter, B.H., Platzbecker, U., Giagounidis, A., Selleslag, D., Labar, B., Germing, U., Salih, H.R., et al. (2011). Low-dose decitabine versus best supportive care in elderly patients with intermediate- or high-risk myelodysplastic syndrome (MDS) ineligible for intensive chemotherapy: final results of the randomized phase III study of the European Organisation for Research and Treatment of Cancer Leukemia Group and the German MDS Study Group. Journal of clinical oncology : official journal of the American Society of Clinical Oncology 29, 1987–1996.

Luo, H., Wei, W., Ye, Z., Zheng, J., and Xu, R.H. (2021). Liquid Biopsy of Methylation Biomarkers in Cell-Free DNA. Trends Mol Med 27, 482–500.

Manara, M.C., Valente, S., Cristalli, C., Nicoletti, G., Landuzzi, L., Zwergel, C., Mazzone, R., Stazi, G., Arimondo, P.B., Pasello, M., et al. (2018). A Quinoline-Based DNA Methyltransferase Inhibitor as a Possible Adjuvant in Osteosarcoma Therapy. Molecular cancer therapeutics 17, 1881–1892.

Markou, A., Londra, D., Tserpeli, V., Kollias, I., Tsaroucha, E., Vamvakaris, I., Potaris, K., Pateras, I., Kotsakis, A., Georgoulias, V., et al. (2022). DNA methylation analysis of tumor suppressor genes in liquid biopsy components of early stage NSCLC: a promising tool for early detection. Clinical epigenetics 14, 61.

Merlo, A., Herman, J.G., Mao, L., Lee, D.J., Gabrielson, E., Burger, P.C., Baylin, S.B., and Sidransky, D. (1995). 5’ CpG island methylation is associated with transcriptional silencing of the tumour suppressor p16/CDKN2/MTS1 in human cancers. Nature medicine 1, 686–692.

Oki, Y., Jelinek, J., Shen, L., Kantarjian, H.M., and Issa, J.P. (2008). Induction of hypomethylation and molecular response after decitabine therapy in patients with chronic myelomonocytic leukemia. Blood 111, 2382–2384.

Oliveira, P.H., Ribis, J.W., Garrett, E.M., Trzilova, D., Kim, A., Sekulovic, O., Mead, E.A., Pak, T., Zhu, S., Deikus, G., et al. (2020). Epigenomic characterization of Clostridioides difficile finds a conserved DNA methyltransferase that mediates sporulation and pathogenesis. Nat Microbiol 5, 166–180.

Otwinowski, Z., Borek, D., Majewski, W., and Minor, W. (2003). Multiparametric scaling of diffraction intensities. Acta Crystallogr A 59, 228–234.

Pappalardi, M.B., Keenan, K., Cockerill, M., Kellner, W.A., Stowell, A., Sherk, C., Wong, K., Pathuri, S., Briand, J., Steidel, M., et al. (2021). Discovery of a first-in-class reversible DNMT1-selective inhibitor with improved tolerability and efficacy in acute myeloid leukemia. Nat Cancer 2, 1002–1017.

Read, R.J., Adams, P.D., Arendall, W.B., 3rd, Brunger, A.T., Emsley, P., Joosten, R.P., Kleywegt, G.J., Krissinel, E.B., Lutteke, T., Otwinowski, Z., et al. (2011). A new generation of crystallographic validation tools for the protein data bank. Structure 19, 1395–1412.

Reich, N.O., Dang, E., Kurnik, M., Pathuri, S., and Woodcock, C.B. (2018). The highly specific, cell cycle-regulated methyltransferase from Caulobacter crescentus relies on a novel DNA recognition mechanism. J Biol Chem 293, 19038–19046.

Ren, R., Horton, J.R., Hong, S., and Cheng, X. (2022). Recent Advances on DNA Base Flipping: A General Mechanism for Writing, Reading, and Erasing DNA Modifications. Advances in experimental medicine and biology 1389, 295–315.

Roberts, R.J., and Cheng, X. (1998). Base flipping. Annu Rev Biochem 67, 181–198.

Rubin, S.J., Hallahan, D.E., Ashman, C.R., Brachman, D.G., Beckett, M.A., Virudachalam, S., Yandell, D.W., and Weichselbaum, R.R. (1991). Two prostate carcinoma cell lines demonstrate abnormalities in tumor suppressor genes. J Surg Oncol 46, 31–36.

Sakai, Y., Suetake, I., Shinozaki, F., Yamashina, S., and Tajima, S. (2004). Co-expression of de novo DNA methyltransferases Dnmt3a2 and Dnmt3L in gonocytes of mouse embryos. Gene Expr Patterns 5, 231–237.

Satange, R., Chuang, C.Y., Neidle, S., and Hou, M.H. (2019). Polymorphic G:G mismatches act as hotspots for inducing right-handed Z DNA by DNA intercalation. Nucleic Acids Res 47, 8899–8912.

Sato, T., Issa, J.J., and Kropf, P. (2017). DNA Hypomethylating Drugs in Cancer Therapy. Cold Spring Harb Perspect Med 7, a026948.

Satpathi, S., Endoh, T., Podbevsek, P., Plavec, J., and Sugimoto, N. (2021). Transcriptome screening followed by integrated physicochemical and structural analyses for investigating RNA-mediated berberine activity. Nucleic Acids Res 49, 8449–8461.

Schuebel, K.E., Chen, W., Cope, L., Glockner, S.C., Suzuki, H., Yi, J.M., Chan, T.A., Van Neste, L., Van Criekinge, W., van den Bosch, S., et al. (2007). Comparing the DNA hypermethylome with gene mutations in human colorectal cancer. PLoS Genet 3, 1709–1723.

Siliciano, J.D., Canman, C.E., Taya, Y., Sakaguchi, K., Appella, E., and Kastan, M.B. (1997). DNA damage induces phosphorylation of the amino terminus of p53. Genes Dev 11, 3471–3481.

Silverman, L.R., Demakos, E.P., Peterson, B.L., Kornblith, A.B., Holland, J.C., Odchimar-Reissig, R., Stone, R.M., Nelson, D., Powell, B.L., DeCastro, C.M., et al. (2002). Randomized controlled trial of azacitidine in patients with the myelodysplastic syndrome: a study of the cancer and leukemia group B. Journal of clinical oncology : official journal of the American Society of Clinical Oncology 20, 2429–2440.

Slupphaug, G., Mol, C.D., Kavli, B., Arvai, A.S., Krokan, H.E., and Tainer, J.A. (1996). A nucleotide-flipping mechanism from the structure of human uracil-DNA glycosylase bound to DNA. Nature 384, 87–92.

Soni, A., Khurana, P., Singh, T., and Jayaram, B. (2017). A DNA intercalation methodology for an efficient prediction of ligand binding pose and energetics. Bioinformatics 33, 1488–1496.

Steffens Reinhardt, L., Groen, K., Newton, C., and Avery-Kiejda, K.A. (2023). The role of truncated p53 isoforms in the DNA damage response. Biochim Biophys Acta Rev Cancer 1878, 188882.

Stephens, C., Reisenauer, A., Wright, R., and Shapiro, L. (1996). A cell cycle-regulated bacterial DNA methyltransferase is essential for viability. Proc Natl Acad Sci U S A 93, 1210–1214.

Stomper, J., Ihorst, G., Suciu, S., Sander, P.N., Becker, H., Wijermans, P.W., Plass, C., Weichenhan, D., Bisse, E., Claus, R., et al. (2019). Fetal hemoglobin induction during decitabine treatment of elderly patients with high-risk myelodysplastic syndrome or acute myeloid leukemia: a potential dynamic biomarker of outcome. Haematologica 104, 59–69.

Stomper, J., Rotondo, J.C., Greve, G., and Lubbert, M. (2021). Hypomethylating agents (HMA) for the treatment of acute myeloid leukemia and myelodysplastic syndromes: mechanisms of resistance and novel HMA-based therapies. Leukemia 35, 1873–1889.

Tian, L., Kim, M.S., Li, H., Wang, J., and Yang, W. (2018). Structure of HIV-1 reverse transcriptase cleaving RNA in an RNA/DNA hybrid. Proc Natl Acad Sci U S A 115, 507–512.

Tsegay, P.S., Hernandez, D., Qu, F., Olatunji, M., Mamun, Y., Chapagain, P., and Liu, Y. (2023). RNA-guided DNA base damage repair via DNA polymerase-mediated nick translation. Nucleic Acids Res 51, 166–181.

Tsumura, A., Hayakawa, T., Kumaki, Y., Takebayashi, S., Sakaue, M., Matsuoka, C., Shimotohno, K., Ishikawa, F., Li, E., Ueda, H.R., et al. (2006). Maintenance of self-renewal ability of mouse embryonic stem cells in the absence of DNA methyltransferases Dnmt1, Dnmt3a and Dnmt3b. Genes Cells 11, 805–814.

Valente, S., Liu, Y., Schnekenburger, M., Zwergel, C., Cosconati, S., Gros, C., Tardugno, M., Labella, D., Florean, C., Minden, S., et al. (2014). Selective non-nucleoside inhibitors of human DNA methyltransferases active in cancer including in cancer stem cells. J Med Chem 57, 701–713.

van der Ploeg, L.H., and Flavell, R.A. (1980). DNA methylation in the human gamma delta beta-globin locus in erythroid and nonerythroid tissues. Cell 19, 947–958.

Veland, N., Lu, Y., Hardikar, S., Gaddis, S., Zeng, Y., Liu, B., Estecio, M.R., Takata, Y., Lin, K., Tomida, M.W., et al. (2019). DNMT3L facilitates DNA methylation partly by maintaining DNMT3A stability in mouse embryonic stem cells. Nucleic Acids Res 47, 152–167.

Wang, L.H., Wu, C.F., Rajasekaran, N., and Shin, Y.K. (2018). Loss of Tumor Suppressor Gene Function in Human Cancer: An Overview. Cell Physiol Biochem 51, 2647–2693.

Wilson, D.M., Duncton, M.A.J., Chang, C., Lee Luo, C., Georgiadis, T.M., Pellicena, P., Deacon, A.M., Gao, Y., and Das, D. (2021). Early Drug Discovery and Development of Novel Cancer Therapeutics Targeting DNA Polymerase Eta (POLH). Front Oncol 11, 778925.

Woodcock, C.B., Horton, J.R., Zhang, X., Blumenthal, R.M., and Cheng, X. (2020). Beta class amino methyltransferases from bacteria to humans: evolution and structural consequences. Nucleic Acids Res 48, 10034–10044.

Woodcock, C.B., Yakubov, A.B., and Reich, N.O. (2017). Caulobacter crescentus Cell Cycle-Regulated DNA Methyltransferase Uses a Novel Mechanism for Substrate Recognition. Biochemistry 56, 3913–3922.

Yoo, C.B., and Jones, P.A. (2006). Epigenetic therapy of cancer: past, present and future. Nat Rev Drug Discov 5, 37–50.

Zeng, Y., Ren, R., Kaur, G., Hardikar, S., Ying, Z., Babcock, L., Gupta, E., Zhang, X., Chen, T., and Cheng, X. (2020). The inactive Dnmt3b3 isoform preferentially enhances Dnmt3b-mediated DNA methylation. Genes Dev 34, 1546–1558.

Zhou, J., Horton, J.R., Blumenthal, R.M., Zhang, X., and Cheng, X. (2021). Clostridioides difficile specific DNA adenine methyltransferase CamA squeezes and flips adenine out of DNA helix. Nat Commun 12, 3436.

Zhou, J., Horton, J.R., Menna, M., Fiorentino, F., Ren, R., Yu, D., Hajian, T., Vedadi, M., Mazzoccanti, G., Ciogli, A., et al. (2023). Systematic Design of Adenosine Analogs as Inhibitors of a Clostridioides difficile-Specific DNA Adenine Methyltransferase Required for Normal Sporulation and Persistence. J Med Chem 66, 934–950.

Zwergel, C., Schnekenburger, M., Sarno, F., Battistelli, C., Manara, M.C., Stazi, G., Mazzone, R., Fioravanti, R., Gros, C., Ausseil, F., et al. (2019). Identification of a novel quinoline-based DNA demethylating compound highly potent in cancer cells. Clinical epigenetics 11, 68.

